# Dynamics of Prion Proliferation Under Combined Treatment of Pharmacological Chaperones and Interferons via a Mathematical Model

**DOI:** 10.1101/2020.07.06.190637

**Authors:** Doménica N. Garzón, Yair Castillo, M. Gabriela Navas-Zuloaga, Nora Culik, Leah Darwin, Abigail Hardin, Anji Yang, Carlos Castillo-Garsow, Karen Ríos-Soto, Leon Arriola, Aditi Ghosh

## Abstract

Prion diseases are lethal neurodegenerative disorders such as mad cow disease in bovines, chronic wasting disease in cervids, and Creutzfeldt-Jakob disease in humans. They are caused when the prion protein PrP^C^ misfolds into PrP^Sc^, which is capable of inducing further misfolding in healthy PrP^C^ proteins. Recent *in vivo* experiments show that pharmacological chaperones can temporarily prevent this conversion by binding to PrP^C^ molecules, and thus constitute a possible treatment. A second strategic approach uses interferons to decrease the concentration of PrP^Sc^. In order to study the quantitative effects of these treatments on prion proliferation, we develop a model using a non-linear system of ordinary differential equations. By evaluating their efficacy and potency, we find that interferons act at lower doses and achieve greater prion decay rates. However, there are benefits in combining them with pharmacological chaperones in a two-fold therapy. This research is crucial to guide future prion experiments and inform potential treatment protocols.

## 1. Introduction

Prions cause fatal diseases that cause irreversible neurodegeneration in the brain. Once infected by prion disease, a person has months, years, or even decades of feeling normal before symptoms appear. Once symptoms begin, the brain slowly becomes spongy, deteriorating where the prions accumulate [1]. This neurodegeneration causes a host of crippling symptoms, like dementia, uncontrollable spasmodic movements (present in Creutzfeldt-Jakob disease), or the inability to sleep (as in fatal familial insomnia). While individuals can be infected by outside sources, such as contaminated meat in the case of mad cow disease, prion diseases can occur spontaneously [2]. Several of these fatal prion diseases are scrapie in sheep; mad cow disease in bovines; chronic wasting disease in cervids and kuru, and Creutzfeldt-Jakob disease in humans [2]. These diseases affect the brain, causing neurons to die; this eventually leads to the death of the individual, as the brain cannot perform its essential functions [2]. Currently, prion diseases have no cure [3], so any strides towards a treatment are important. Even though prion diseases are far from commonplace, they are fatal and kill hundreds of people every year. In 2017 alone, over 500 people in the United States died from Creutzfeldt-Jakob disease [4]. Further, the study of prion diseases has implications for other neurodegenerative diseases, such as Alzheimer’s and Parkinson’s, as these illnesses are very similar to prion diseases. That is, they involve the loss of function of the PrP protein which negatively affects the brain’s function [5].

Little is known about the specific functions of the prion protein, PrP; however, it is known that PrP slows neuronal apoptosis (cell death) [6]. Prions are created when the protease resistant protein (PrP) misfolds. These proteins appear normally in mammalian brains. The mechanisms of this folding error are not yet fully understood. However, it is known that misfolded proteins can cause other properly folded PrP to form into a prion. This correctly folded form is sometimes called PrPC (C for “cellular”). Problems begin when this protein folds into an isoform^1^, called PrP^Sc^ (Sc stands for “scrapie”). Prions have no DNA or RNA themselves, so they go against the central dogma of biology because they are still able to replicate by inducing further misfolding [1]. The “protein only hypothesis” proposes that prion replication happens without the involvement of nucleic acid [1]. When PrP folds normally into PrP^C^, its folded form is rich in *α*-helices. If PrP folds into a *β*-sheet-rich form instead of one rich in *α*-helix structures, it forms PrP^Sc^ and thus becomes a prion [8].

There are two common hypotheses used to describe prion spread. The first hypothesis is the heterodimer^2^ model. This model assumes that when a PrP^Sc^ protein comes into contact with a PrP^C^ protein, the prion unfolds the healthy protein and acts as a template to turn the PrPC into PrP^Sc^ (see Figure 1) [10]. This simple model, however, does not include the experimental fact that prions form polymers: chains of PrP^Sc^ monomers. The second hypothesis is called the nucleated polymerization model, and it studies chains of prions and how the chain length varies [11]. When a chain of prions infects a new monomer, it adds the PrPC to the chain, causing it to misfold, and the chain grows by one monomer. However, if the chain breaks, there are two options. First, the polymer can break into two smaller polymers; second, if one of the polymer’s length is below a certain threshold, it dissociates into monomers, [12]. In this model, the monomers are not infectious by themselves, but they can join an existing polymer. This work will take into consideration the polymerization model (see Figure 2). These replication models are usually in one or two spatial dimensions. The two-dimensional models study prion aggregations, and they have explained how incubation times and inoculation doses are highly correlated [13]. Essentially, the period before the symptoms appear is related to the amount of prions going into the brain [13]. The one-dimensional models treat PrP^Sc^ as fibrillic structures that can add new monomers on either end of a chain. This has not only been experimentally studied [14] but also it has been widely analyzed mathematically [12, 15, 16, 17].

**Figure 1:**
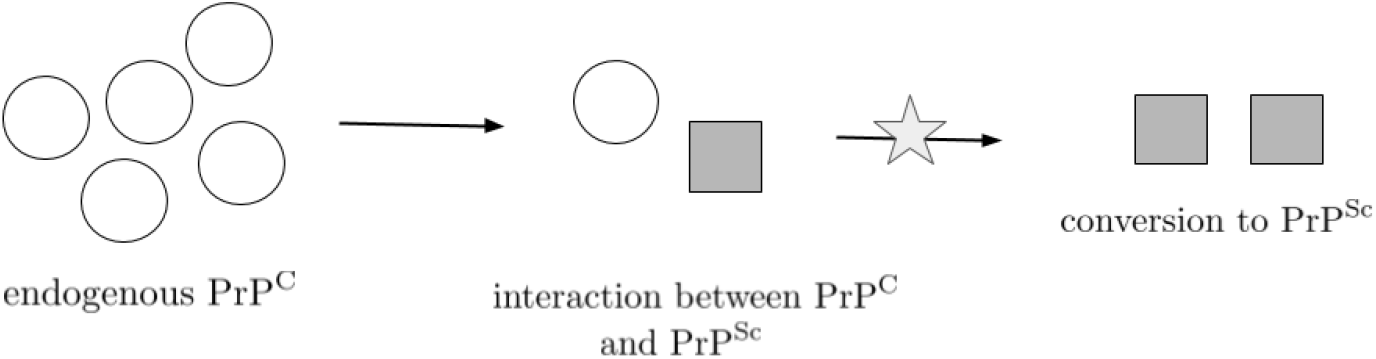
The Heterodimer model shows how one monomer of misfolded protein, PrP^Sc^, converts a monomer of PrP^C^ [9].

**Figure 2:**
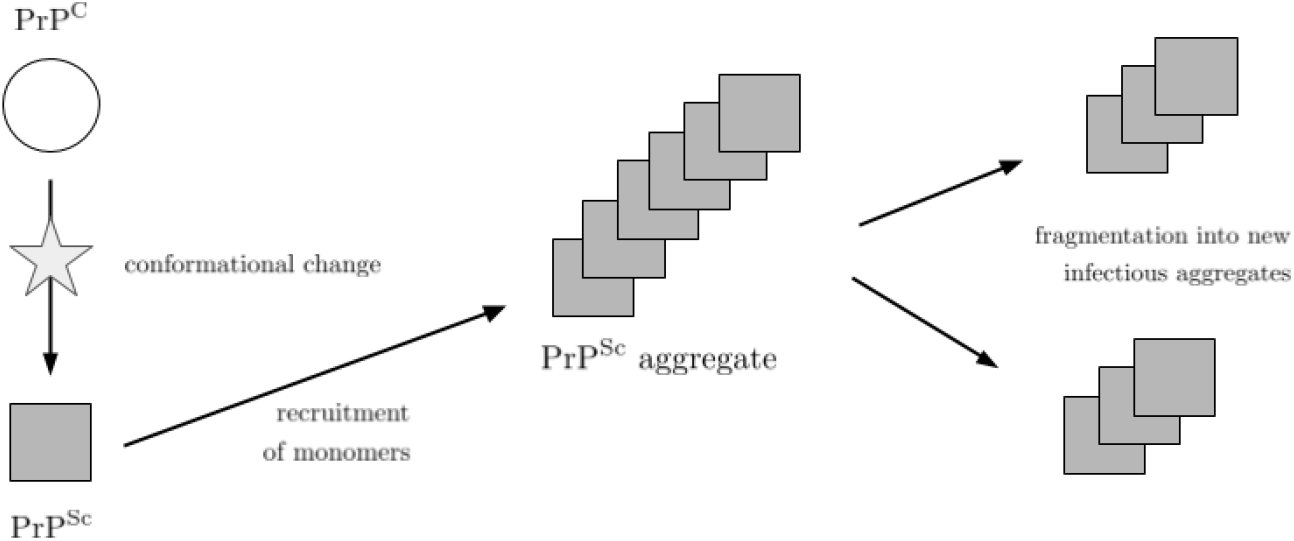
The polymerization model describes a prion proliferation hypothesis in which PrP^Sc^ monomers form polymer chains. These aggregates are infectious and actively recruit free endogenous PrP^C^ protein monomers that have undergone conformational change into PrP^Sc^ monomers. Large PrP^Sc^ polymer chains eventually become unstable and break into new infectious units and repeat the process [10].

### 1.1. Possible Treatments for Prion Diseases

Recent experimental research has shown that there are possible treatments for prion diseases [18, 19, 20, 8]. These treatments can be categorized in four mechanistic ways, as described by Kamatari [21]. The first mechanism (I) is a stabilization of the PrPC structure by direct association of a molecule to prevent formation to the PrP^Sc^ isoform. Mechanism II is an indirect association between the interfering molecule and PrPC; this blocks the interaction between PrPC and PrP^Sc^, slowing the rate at which PrP^Sc^ is able to spread. Mechanism III removes PrPC from the system, and mechanism IV prevents PrP^Sc^ from proliferating by associating PrP^Sc^ with molecules other than PrPC.

This paper will focus on Mechanism I, specifically through the use of pharmacological chaperones^3^. Many different types of molecules are able to act as pharmacological chaperones. It has been shown that antibodies are able to act mechanistically as pharmacological chaperones do [23].

*In vivo* experiments have shown that antibodies can be used to block the proliferation of prions [18] by forcing the secondary structure of PrP protein into an α-helix form rather than *β*-sheets which are associated with PrP^Sc^ (see Figure 3). Specially-engineered molecules can be designed to bind at locations critical to the correct folding of PrP^C^ [24]. However, pharmacological chaperones tend to have a short half-life (though the specific half-life depends on the drug that is being used), which means that eventually treated PrP^C^ will become susceptible to misfolding again [25]. This suggests that we must keep the concentration of pharmacological chaperones high in order to keep the disease at bay.

**Figure 3:**
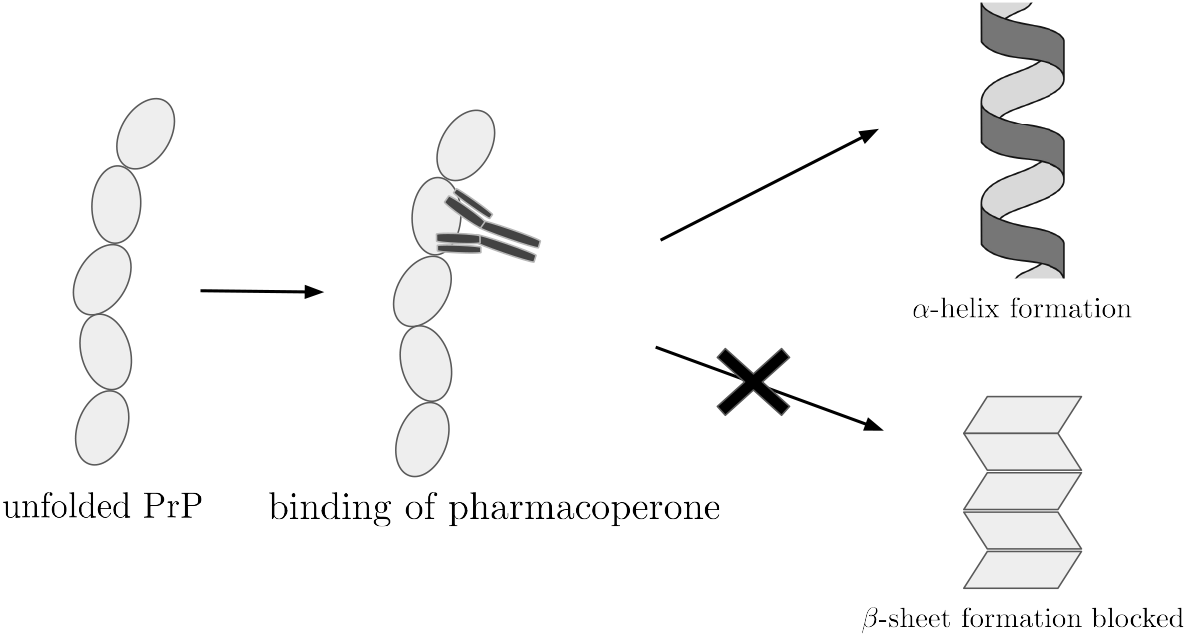
Pharmacological chaperones bind to unfolded PrP, forcing it to fold into *α*-helix-rich form and thus PrP^C^ instead of PrP^Sc^ which is rich in *β*-sheets.

Another treatment involves interferons, a part of the immune system that raises the body’s immune response by signaling other proteins [26]. It has been documented that there is a naturally occurring increase in type I interferon (I-IFN) expression in a brain affected with a prion disease. This has been shown to be the case in scrapie-infected hamster and mouse brains by identifying upregulated gene expression [27, 28]. However, it is not well-known how this innate reaction recognizes and attacks the infected proteins given that PrP^C^ and PrP^Sc^ do not differ in amino acid sequences [1]. Ishibashi et al. (2012) [29] proposed that PrP^Sc^ infection induces a response from Toll-like receptor (TLR) proteins, a class of pattern-recognizing receptors essential to the immune response signalling pathway [30]. These proteins are upstream of I-IFN, which results in a cascade of signals inducing immune response [29]. Moreover, *ex vivo* experiments in scrapie infected mice have shown that inducing I-IFN via TLR signalling reduces PrP^Sc^ concentration in the model host during the early stages of infection [31]. With these two treatments, prion formation can be slown with pharmacological chaperones and the prion population can be significantly diminished with interferons.

In this study, a possible treatment for prion diseases is considered: a twofold therapy, a combination of pharmacological chaperones and interferons. The findings of this research evaluates the efficacy and potency of these treatments within their safe ranges. As both pharmacological chaperones and interferons do reduce the concentration of prions, we examine whether or not a combination of sub-maximal dosages would work. This work has implications for prion disease therapies, diseases which are currently not only incurable but untreatable.

This paper constructs a non-linear system of differential equations based on the previous work by Nowak et al. [12] with the addition of interferon and pharmacological chaperone treatments. We find the system’s equilibrium points and examine their stability, as well as analyze important indicators such as R_0_ (basic reproduction number) and the growth rate of prion proliferation. Numerical simulations provide even more insight into each treatment on its own, as well as their combination. We conclude with an examination of efficacy and potency and answer the question of how these treatments can be used to treat prion diseases.

This model implements the polymerisation hypothesis for proliferation of prions. Two types of treatments are incorporated in this biological model. The first one consists of a dose of pharmacological chaperones which prevent prion formation based on *in vivo* experiments by Gunther et al. [32]; the second treatment consists of a dose of interferons that decreases the amount of prions in the brain [31]. Our model is based on two previous mathematical models. Masel et al. (1999) [10] used the hypothesis of nucleated polymerisation as the mechanism of proliferation. Masel et al. then established a deterministic infinite dimensional dynamical system to model the dynamics between the population of the susceptible monomers and the polymers of prions with a distinct equation to describe the population of polymers of each possible length; however, this paper does not consider any treatments. In a subsequent paper, Masel et al. (2000) [15] used a theoretical kinetic model to calculate the growth rate of protein aggregates as a function of certain drugs which blocks the ends of amyloids. However, the treatment examined in that model differs mechanistically from those examined here.

Figure 6 describes the kinetic model in detail. To consider the dynamics of this system, the model posed here examines what can happen with a PrP^C^ monomer. PrPC monomers are naturally produced at a rate Λ. There are several paths that this monomer can take once it has joined the system. A polymer of PrP^Sc^ can convert this monomer, which happens at rate *β_S_*. From there, the polymer will either split or simply grow longer. The breakage rate is *b_i,j_*, where *i* is the length of the polymer and *j* (and thus *i* – *j*) are the lengths of the new polymers after it splits. When the polymer breaks, two events can happen. It can break in two smaller polymers, each with a length greater than *n*, and they will continue converting monomers. Otherwise, it can split into a polymer and small chain (whose length is less than the threshold *n*) which will dissociate into separate PrP^Sc^ monomers (see Figure 7 for a visual depiction of the breakage process). This monomer can join an existing polymer at rate *β_R_*, but it cannot be treated with pharmacological chaperones to prevent it from rejoining a chain as the pharmacological chaperones can only act on PrP^C^. The introduction of interferons, however, may mean that our polymer is eliminated much sooner. The interferons induce an additional death rate for PrP^Sc^, called *μ_I_*. If the PrP^C^ monomer is not treated with pharmacological chaperones, it can be infected. The addition of pharmacological chaperones into the system can be described by the dosage rate, *D*. Once a PrPC monomer has been treated, it will be immune from PrP^Sc^ until either the pharmacological chaperone or the PrP^C^ monomer degrades. Degradation of pharmacological chaperones and PrPC happen at rates *μ_A_* and *μ_S_*, respectively. These interactions are summarized mathematically in the equations section.

**Figure 4:**
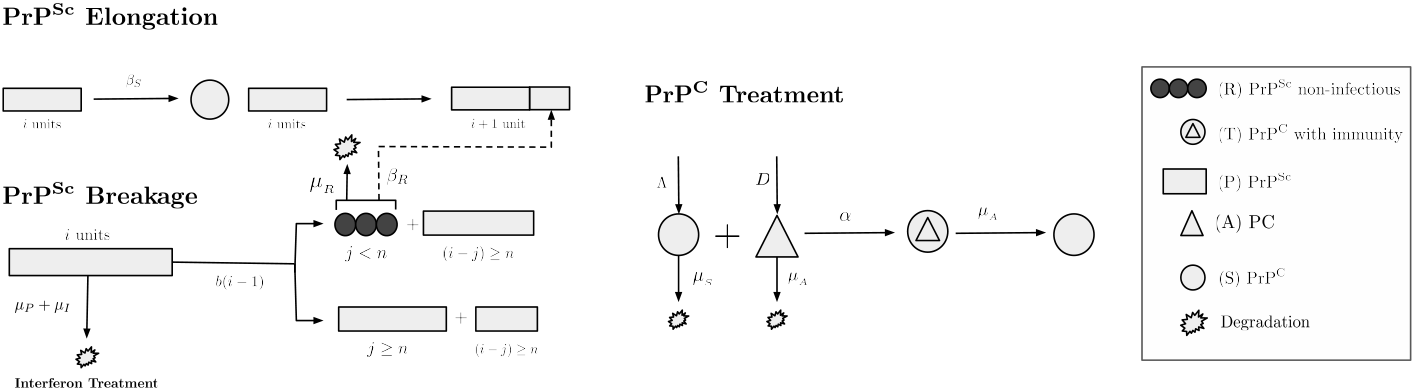
Detailed kinetic model that includes the nucleated polymerisation hypothesis and the incorporation of the treatments.

**Figure 5:**
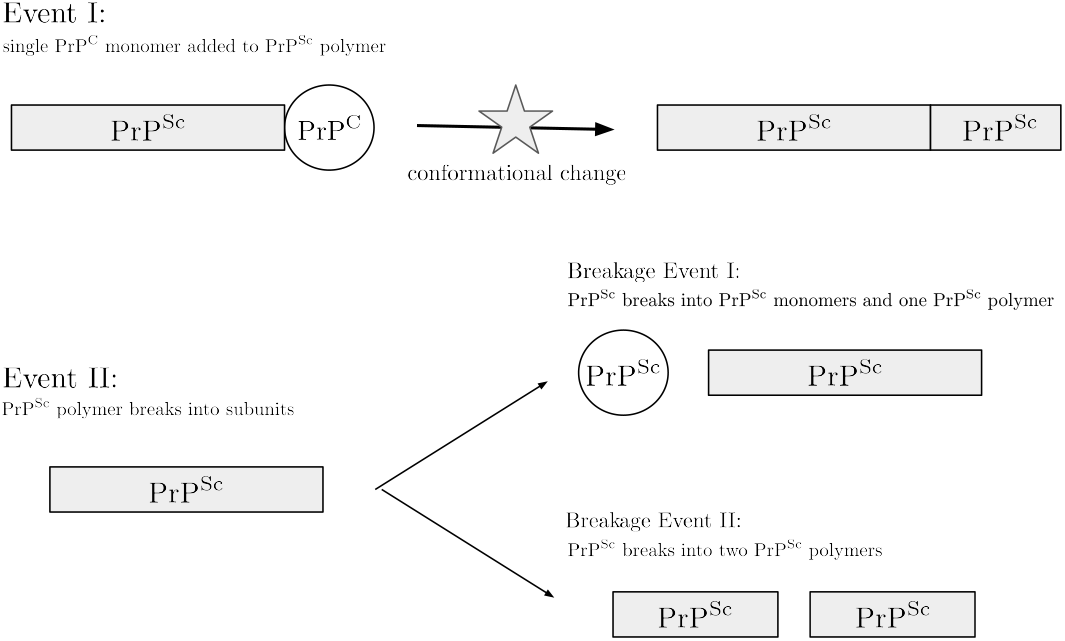
The possible events for a PrP^Sc^ polymer. In event I, a PrP^Sc^ polymer contacts a PrP^C^, inducing a conformational change in which the PrP^C^ is added to the polymer. In event II, the PrP^Sc^ breaks into fragments. There are two unique breakage events. In breakage event I the PrP^Sc^ polymer breaks to create one PrP^Sc^ monomer and one PrP^Sc^ polymer. In breakage event II, the PrP^Sc^ polymer breaks to create two PrP^Sc^ polymers.

**Figure 6:**
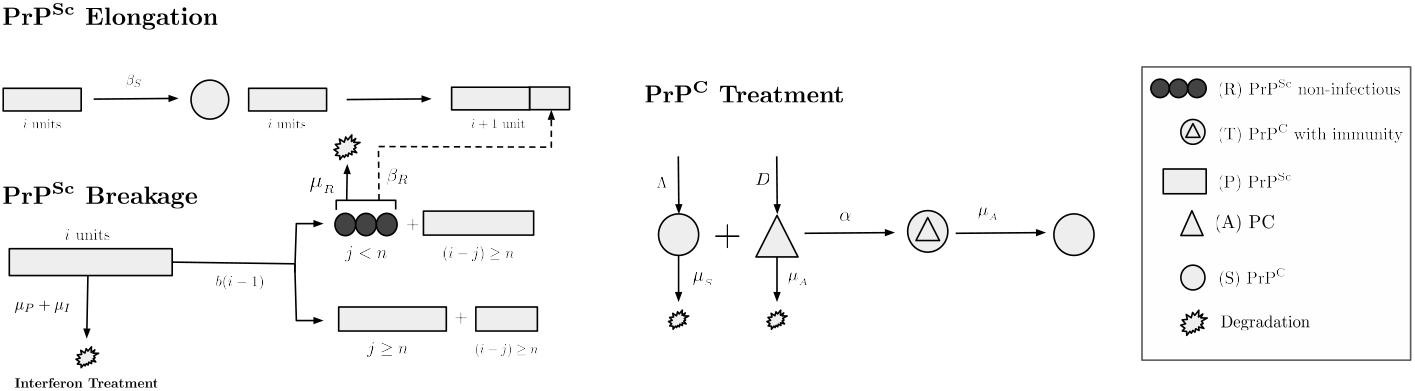
Detailed kinetic model that includes the nucleated polymerisation hypothesis and the incorporation of the treatments.

**Figure 7:**
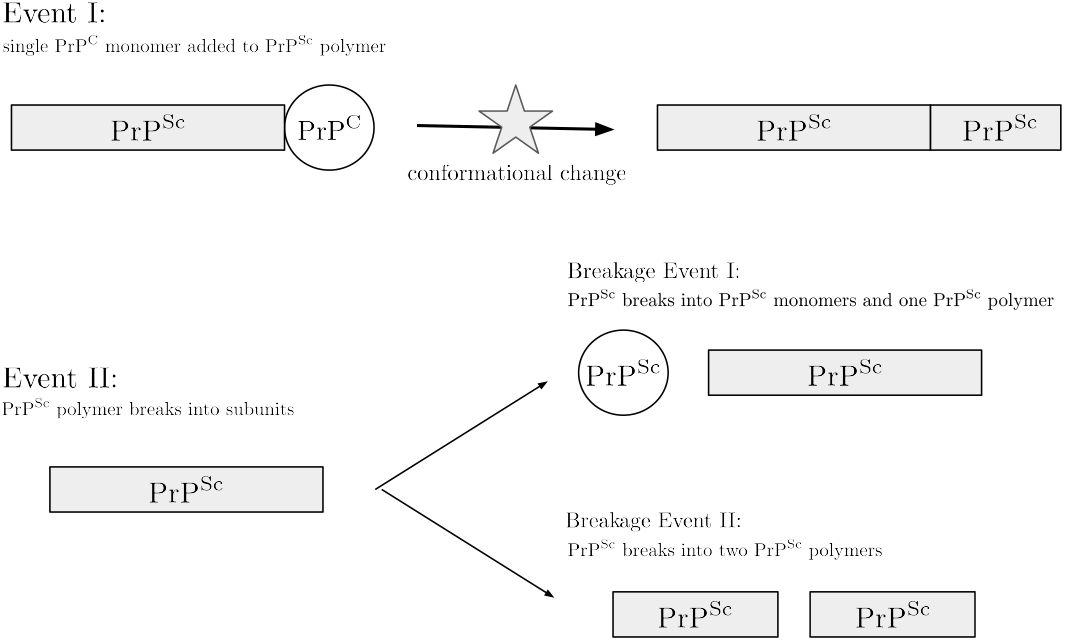
The possible events for a PrP^Sc^ polymer. In event I, a PrP^Sc^ polymer contacts a PrP^C^, inducing a conformational change in which the PrP^C^ is added to the polymer. In event II, the PrP^Sc^ breaks into fragments. There are two unique breakage events. In breakage event I the PrP^Sc^ polymer breaks to create one PrP^Sc^ monomer and one PrP^Sc^ polymer. In breakage event II, the PrP^Sc^ polymer breaks to create two PrP^Sc^ polymers.

## 2. Prion Proliferation Treatment Model

This model implements the polymerisation hypothesis for proliferation of prions. Two types of treatments are incorporated in this biological model. The first one consists of a dose of pharmacological chaperones which prevent prion formation based on *in vivo* experiments by Gunther et al. [32]; the second treatment consists of a dose of interferons that decreases the amount of prions in the brain [31]. Our model is based on two previous mathematical models. Masel et al. (1999) [10] uses the hypothesis of nucleated polymerisation as the mechanism of proliferation. Masel et al. then establishes a deterministic infinite dimensional dynamical system to model the dynamics between the population of the susceptible monomers and the polymers of prions with a distinct equation to describe the population of polymers of each possible length; however, this paper does not consider any treatments. In a subsequent paper, Masel et al. (2000) [15] uses a theoretical kinetic model to calculate the growth rate of protein aggregates as a function of certain drugs which blocks the ends of amyloids. However, the treatment examined in that model differs mechanistically from those examined here.

Figure 6 describes the kinetic model in detail. To consider the dynamics of this system, the model posed here examines what can happen with a PrP^C^ monomer. PrP^C^ monomers are naturally produced at a rate Λ. There are several paths that this monomer can take once it has joined the system. A polymer of PrP^Sc^ can convert this monomer, which happens at rate *β_S_*. From there, the polymer will either split or simply grow longer. The breakage rate is *b_i,j_*, where *i* is the length of the polymer and *j* (and thus *i* – *j*) are the lengths of the new polymers after it splits. When the polymer breaks, two events can happen. It can break in two smaller polymers, each with a length greater than *n*, and they will continue converting monomers. Otherwise, it can split into a polymer and small chain (whose length is less than the threshold *n*) which will dissociate into separate PrP^Sc^ monomers (see Figure 7 for a visual depiction of the breakage process). This monomer can join an existing polymer at rate *β_R_*, but it cannot be treated with pharmacological chaperones to prevent it from rejoining a chain as the pharmacological chaperones can only act on PrPC. The introduction of interferons, however, may mean that our polymer is eliminated much sooner. The interferons induce an additional death rate for PrP^Sc^, called *μ_I_*. If the PrP^C^ monomer is not treated with pharmacological chaperones, it can be infected. The addition of pharmacological chaperones into the system can be described by the dosage rate, *D*. Once a PrPC monomer has been treated, it will be immune from PrP^Sc^ until either the pharmacological chaperone or the PrP^C^ monomer degrades. Degradation of pharmacological chaperones and PrPC happen at rates *μ_A_* and *μ_S_*, respectively. These interactions are summarized mathematically in the equations section.

The basic assumptions for this model come from the published study by Masel et al. [10]. For instance, the rate at which PrPC and non-infectious PrP^Sc^ molecules are converted into infectious PrP^Sc^ polymers is the same (*β_S_* = *β_R_*), and is independent on the length of the PrP^Sc^ chain. Also, the rate at which prions break is the same for all lengths *i* ≥ *n*, so we let *b_i,j_* = *b*. Furthermore, the death rates for PrP^C^ and non-infectious PrP^Sc^ are the same (*μ_S_* = *μ_R_*) [10]. It is important to note that death rate of PrP^Sc^, *μ_P_*, is different. The model allows *μ_I_* to either affect the non-infectious PrP^Sc^ monomers or not; this is achieved by the term *σ*, where 0 ≤ *σ* ≤ 1 is a probability representing the degree to which the PrP^Sc^ monomers are affected by the interferons.

The transmission of prions follows the polymerisation hypothesis. The prion polymers are considered linear, that is, PrP^C^ can only attach to the ends of PrP^Sc^ polymers [10]. For simplicity the population of PrPC is considered to be a well-mixed homogeneous system. Since the reaction that involves the misfolding of a protein are extremely fast and are measured in microseconds [33], we consider the time to fold or unfold a protein is negligible. We incorporate treatments as constant dosages of pharmacological chaperones and interferons over time and make several assumptions about them. For the first treatment, each one of the pharmacological chaperones introduced in the body binds to one PrP^C^ in order to prevent the misfolding process. The pharmacological chaperone in question has high specificity, meaning that we neglect the rate at which this pharmacological chaperone binds to molecules other than PrP^C^. Also, we assume that pharmacological chaperones do not reduce the amount of prions directly, which means that they act as a blocking monomer treatment. Lastly, our model includes no spatial dependence.

In this model, a non-linear system of ordinary differential equations is studied. This model introduces pharmacological chaperones and interferons into a prion-infected brain. The system of infinite differential equations is presented at Appendix, is reduced to a closed system of six equations that will be used to study the dynamics of the prion population in an individual’s brain.

*T* and *A*, measure treated PrPC and pharmacological chaperones respectively, as in system A.1 in Appendix. Instead of an equation for each polymer of length i, we have a class *P*, which counts polymers of PrP^Sc^. *P* is defined as 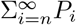. An additional class must also be introduced in order to close this system. We define *Z* to be the total number of PrP^Sc^ monomers in the polymer chains, formally 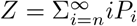.

The closed system of equations is then given by:

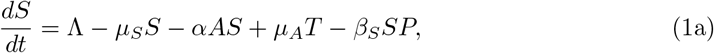

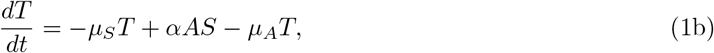

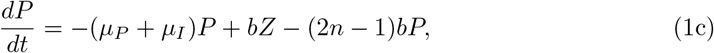

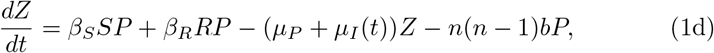

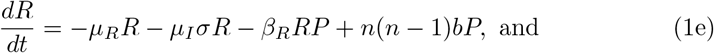

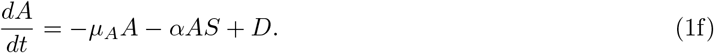

Here, *P* represents the number of PrP^Sc^ polymers and *Z* represents the total number of PrP^Sc^ monomers within those chains. For details on how the system was closed, see Appendix. The parameters used to describe the dynamics of the system are summarized in Table 1.

**Table 1:** Table of parameters for the ODE’s of (1), where 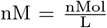

| Name | Description | Value | Units | Reference |
| --- | --- | --- | --- | --- |
| State variables |  | Initial Condition |  |  |
| $S$ | Susceptible population of PrP<sup>C</sup> | [300,750] | nM | [34] |
| $T$ | PrP<sup>C</sup> proteins treated with pharmacological chaperones | 0 | nM | |
| $P$ | PrP<sup>Sc</sup> chains | 3 | nM | |
| $Z$ | Total of monomers for each PrP<sup>Sc</sup> polymer in the chains | 90 | nM | |
| $R$ | PrP<sup>Sc</sup> smaller than $n$ | 0 | nM | |
| $A$ | Pharmacological chaperone population | 0 | nM | |
| Parameters |  |  |  |  |
| $\Lambda$ | Natural production of PrP<sup>C</sup> | [1800, 3000] | $\frac{\text{nM}}{\text{day}}$ | [35] |
| $\mu_S$ | Degradation rate of PrP<sup>C</sup> | [3,5] | $\frac{1}{\text{day}}$ | [35] |
| $\mu_P$ | Degradation rate of PrP<sup>Sc</sup> polymers | [0.02,0.2] | $\frac{1}{\text{day}}$ | [10] |
| $\mu_R$ | Degradation rate of PrP<sup>Sc</sup> monomers | [3,5] | $\frac{1}{\text{day}}$ | [35] |
| $\mu_A$ | Degradation rate of pharmacoperones | 1/29 | $\frac{1}{\text{day}}$ | [36] |
| $\mu_I$ | Degradation rate of PrP<sup>Sc</sup> due to interferons | [0,0.2511] | $\frac{1}{\text{day}}$ | |
| $\sigma$ | Effect of interferons on R degradation | [0,1] | — | |
| $\beta_S$ | Infection rate of PrP<sup>C</sup> | 0.8 | $\frac{1}{(\text{day})\text{nM}}$ | [16] |
| $\beta_R$ | Infection rate of PrP<sup>Sc</sup> monomers | 0.8 | $\frac{1}{(\text{day})\text{nM}}$ | [16] |
| $\alpha$ | Rate that a pharmacoperone binds to a PrP<sup>C</sup> | 46.656 | $\frac{1}{(\text{day})\text{nM}}$ | [37] |
| $b_{i,j}$ | Rate of breakages of the PrP<sup>Sc</sup> of length $i$ in a PrP<sup>Sc</sup> of length $j$ and $i - j$ | 0.00032 | $\frac{1}{\text{day}}$ | [16] |
| $n$ | Minimum polymer length | {2,3,4,5,6} | — | [35, 38] |
| $D$ | Daily dosage of pharmacological chaperones | [0,5816] | $\frac{\text{nM}}{\text{day}}$ | [39] |

### 2.1. Linking interferon dosage to prion mortality

In this model, the interferon treatment is represented by the interferon-induced mortality of PrP^Sc^ chains, *μ_I_*. In order to define the functional form linking *μ_I_* with an interferon daily dosage *I*, we analyzed recent experimental data measuring prion degradation after inoculation with interferons [31]. The study reports the ratio between prion concentrations in treated and untreated mice, 48h after the inoculation with increasing interferon dosages. We quantified the coefficients for the best-fit Hill function interpolating the data and obtained

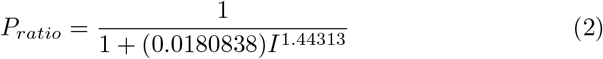

where *I* is the average daily concentration of interferons (for details, see Appendix C).

Assuming that the prion concentration decays exponentially in time and that *μ_I_* can be written as *μ_P_* multiplied by a dose-dependent constant *k*(*I*), we obtain the equation

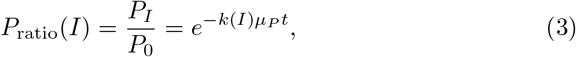

where *t* is the time since inoculation. Because the experiment measured concentrations after 48 hours, we set *t* = 2 days and solve for *k*:

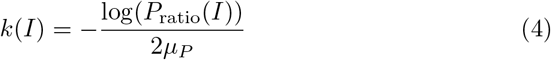

where *P*_ratio_(*I*) is a ratio between 0 and 1 defined by Equation Appendix B.1.2. Thus, in the following sections we consider

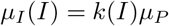

## 3. Analysis

For analysis, the model is examined in several different cases: both treatments are being administered (*D* ≠ 0 and *μ_I_* ≠ 0), only pharmacological chaperones administered (*D* ≠ 0 and *μ_I_* = 0), and only interferons administered (*D* = 0 and *μ_I_* = 0). The no treatments case (*D* = 0 and *μ_I_* = 0) is very similar to the model described in Masel et al. [10]. Any differences in analysis between this case and the one presented by Masel et al. is noted.

### 3.1. Existence of Prion-Free Equilibrium

In order to calculate the prion-free equilibrium (PFE), assume the population of prions *P* is zero. Then from Equations and 1d and 1e of System 1, the following equations are obtained:

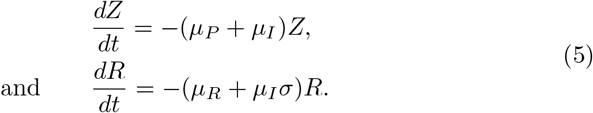

Therefore,

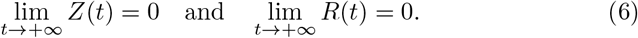

Using equation 1a, *S* satisfies the equation

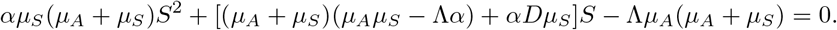

The roots of this quadratic equation are

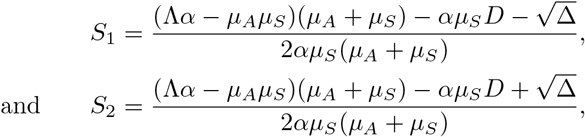

where Δ = [(*μ_A_μ_S_* – Λ_*α*_)(*μ_A_* + *μ_S_*) + *αμ_S_D*]^2^ + 4*α*Λ*μ_S_μ_A_*(*μ_A_* + *μ_S_*)^2^. Notice that Δ > 0, *αμ_S_*(*μ_A_* + *μ_S_*) > 0, –Λ*μ_A_*(*μ_A_* + *μ_S_*) < 0 and *S*_2_ > *S*_1_ (See more details in Appendix #). Then this quadratic equation must have a positive root *S*^*^ = *S*_2_. From equations 1b and 1f, we obtain the relations

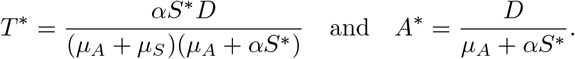

That is, *T*^*^ and *A*^*^ exist and are positive, and they must have biological relevance. Therefore, the prion-free equilibrium must exist.

In our model, the PFE of the system is given by *E*^*^ = (*S*^*^, *T*^*^, 0,0,0, *A*^*^), where

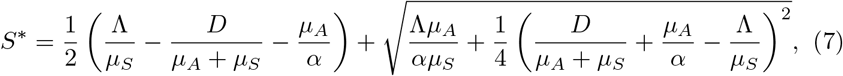

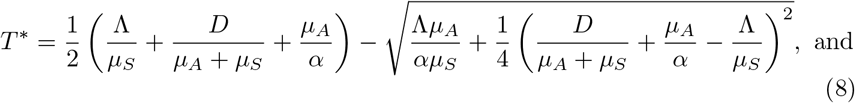

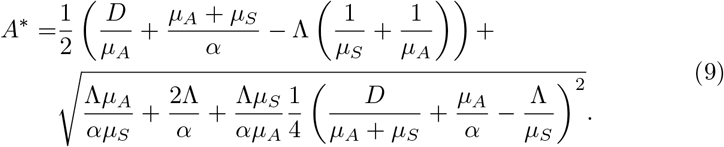

Where *P*^*^ = *Z*^*^ = *R*^*^ = 0 (see Appendix for details). These equilibrium values will always be real and positive, regardless of the parameter values (see Appendix for details).

Additionally, notice that 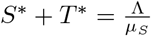, so when *D* = 0 (i.e. when no pharmacological chaperone treatment is being used) the PFE becomes 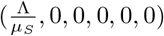, the same equilibrium as in previous models which did not include treatment [10]. It can be seen then that the introduction of the pharmacological chaperone treatment lowers *S*^*^. When *μ_I_* = 0, the prion-free equilibrium does not change with respect to the PFE.

### 3.2. Stability Condition of the Prion-Free Equilibrium

#### Theorem 3.1.

*The model given by System (1a-1f) always has a prion free equilibrium*

*E*^*^ = (*S*^*^, *T*^*^, 0,0,0, *A*^*^) *when R*_0_ < 1, *where*

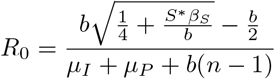

*Moreover, equilibrium E^*^ is locally asymptotically stable when R*_0_ < 1, *and when R*_0_ > 1, *E^*^ is unstable*.

A summary of the proof of the theorem can be seen by examining the eigenvalues of the Jacobian matrix evaluated at *E*^*^, which are the following:

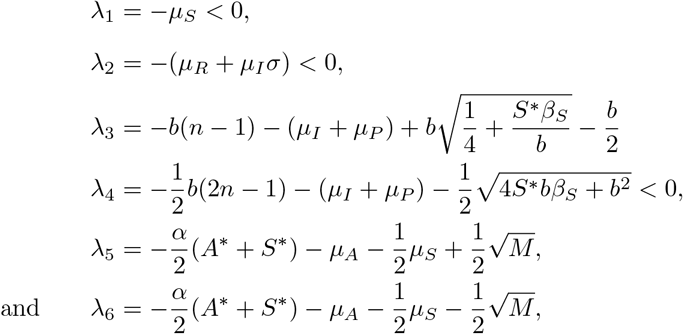

where 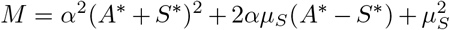. It is easy to show that λ_5_ < 0 and *λ*_6_ < 0 when M is positive. On the other hand, *λ*_3_ < 0 if *ϕ*_1_ < 0 (*R*_0_ < 1), where 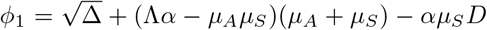, 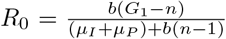. For the details of the full proof, see Appendix. As a result, equilibrium *E*^*^ is locally stable when *ϕ*_1_ < 0 and *R*_0_ < 1, as when those criteria are met all the eigenvalues are negative.

### 3.3. Basic Reproductive Number

In Masel et al. [10], the basic reproductive number (R_0_) of the system was found heuristically, i.e. by multiplying the rate of creation of new prions by the average lifespan of a prion (time spent in *P*). Table 2 shows basic reproductive values found heuristically and through a Next Generation Matrix for both Masel et al.’s system and (1a-1f). Also, values found for system (1a-1f) are written as functions of the treatment paramters *D* and /*mu_I_*, and R_0_ (0, 0) = R_0_ for both the heuristic and Next Generation Matrix values. For either system, when R_0_ = 1 or R_0_(*D, μ_I_*) = 1, the heuristic and Next Generation R_o_ can be reduced to the exact same condition. This implies that these values have the same region of existence [40].

**Table 2:** The R_0_ values for System 2.2 and previous literature found using different methods.

| Case | Heuristic | Next Generation Matrix |
| --- | --- | --- |
| Masel 1999 [10] | $R_0^H = \frac{\sqrt{\beta b S^* + \frac{b^2}{4}} - \frac{b}{2}}{\mu_P + b(n-1)}$ | $R_0^{NG} = \frac{b(\beta S^* - n(n-1)b)}{\mu_P(\mu_P + (2n-1)b)}$ |
| System 2.2 | $R_0^H(D, \mu_I) = \frac{\sqrt{\beta b S^* + \frac{b^2}{4}} - \frac{b}{2}}{(\mu_P + \mu_I) + b(n-1)}$ | $R_0^{NG}(D, \mu_I) = \frac{b(\beta S^* - n(n-1)b)}{(\mu_P + \mu_I)((\mu_P + \mu_I) + (2n-1)b)}$ |

To show the heuristic expressions were determined, take 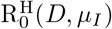. In this model, the terms which represent the degradation of prions are (*μ_P_* + *μ_I_* + (*n* – 1)*b*)*P* and the terms which represent the creation of new prions are *b*(*Z* – *nP*). By normalizing these terms by 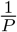, those terms become (*μ_P_* + *μ_I_* + (*n* – 1)*b*) and *b*(*G* – *n*), where 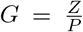 is the average length of prion polymers. The derivative of the average length of the prion polymers (*G*) is

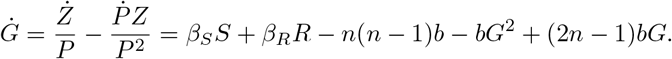

Notice that near the prion-free equilibrium, 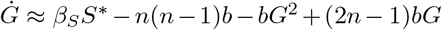. Therefore, this equation approximately describes behavior of a very small initial infection. The roots of this differential equation are

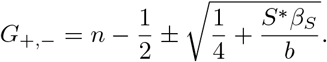

Because *n* is always greater than or equal to one, 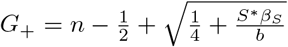 is always positive. The other root, *G*_, will be negative when 0 < *β_s_S*^*^ – *n*(*n* – 1)*b*. This is true when 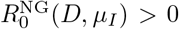, so *G*_ is always negative for biologically-relevant parameter ranges. Thus, as can be seen in the phase diagram below (see Figure 8), once an infection has been introduced to the system, the average polymer length G will reach a fixed value, *G*_+_. That means that the number of secondary infections can be represented by 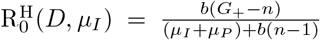. The difference between *n* and *G*_+_ in the numerator represents the minimum size n required to be infectious. Specifically, the smallest possible value for *Z* is *nP*, where each prion chain is of its minimum length. Normalizing each of these terms shows that the minimum size of the average length *G* is *n*, so 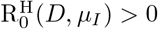 for biologically significant parameter values.

**Figure 8:**
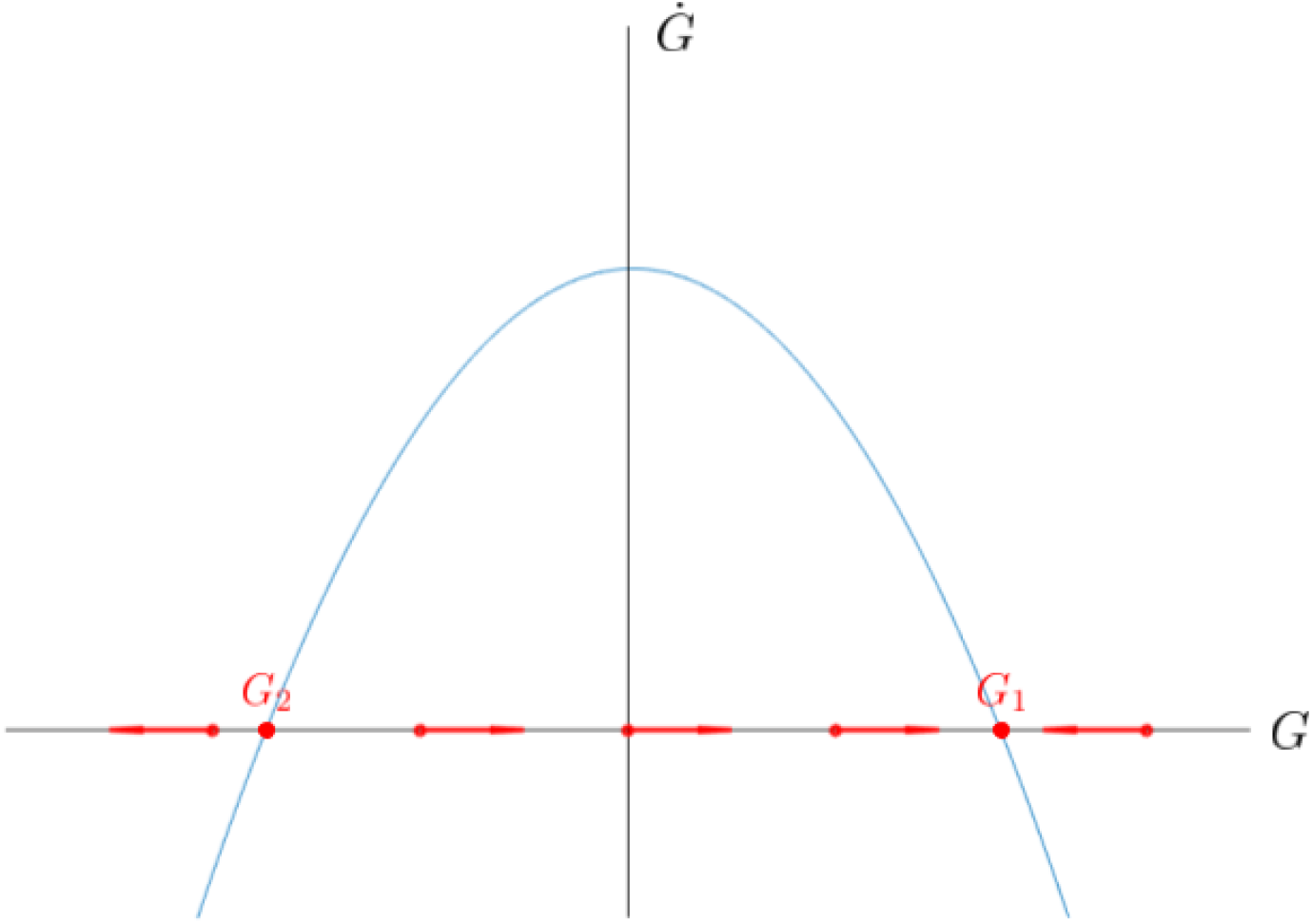
Phase plane for *G*, the average polymer length. *G*_+_ and *G*_–_ are the fixed points of *G. G*_+_ is a stable value for the average polymer length.

Next take the value found using a Next Generation Matrix. The secondary infection rate representing population changes in the number of monomers in prion chains (*Z*) is

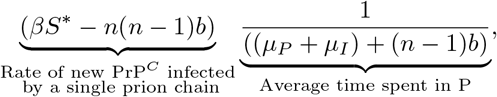

and the condition *βS*^*^ ≫ *b* is required to be true. The secondary infection rate representing population changes in number of infections polymers (*P*) is

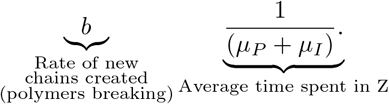

Multiplying these together gives 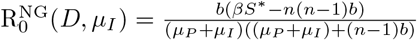. Thus, 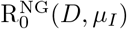 represents prion replication as a two stage process in which a prion must first grow longer and then break in order to create a new infectious chain. Thus, this value represents the secondary infections of *P* over two time steps, rather than one. As such, like many vector disease models, the geometric mean of 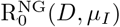 is also a valid basic reproductive number. Interestingly enough, for the parameter values used in this work 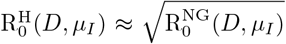 as can be seen in Figure 9.

**Figure 9:**
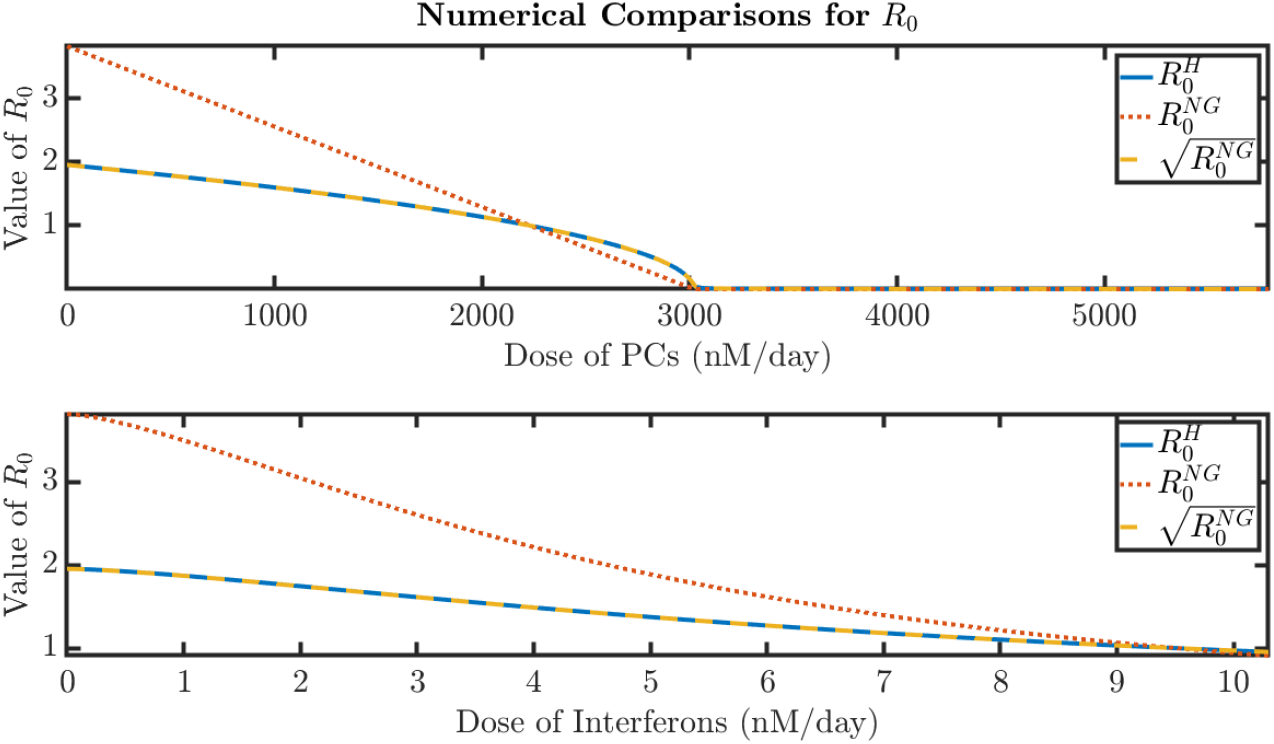
The geometric mean of 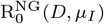 is approximately equal to the heuristic value, 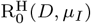. **Top:** Effect of PC dosage *D* on *R*_0_ in the absence of interferons. The parameters are Λ = 3000, *μ_S_* = 5, *μ_R_* = 5, *μ_P_* = 0.2, *n* = 2, *μ_I_* = 0. **Bottom:** Effect of interferon dosage *I* on *R*_0_ in the absence of PC’s. The parameters are Λ = 3000, *μ_S_* = 5, *μ_R_* = 5, *μ_P_* = 0.2, *n* = 2 and *D* = 0

In Figure 9, it is shown the behavior of the basic reproductive number in both cases. First, the dosage of interferons is maintained constant and the basic reproductive number is plotted against the dosage of Pharmacological Chaperons. Here *Ro* becomes less than 1 and it saturates at a certain value, which depends on the value of λ. Second, the basic reproductive number is plotted against the dosage interferons. Here also R_0_ becomes less than 1. Consequently, both treatments are enough to end with the proliferation of prion disease, separately. However, the combination of both treatments is explored in this paper, to understand the benefit of a combined treatment.

When the treatment methods are being implemented, it can be seen that the interferon treatment lowers the R_0_ by increasing the death terms of prion polymers and monomers from *μ_P_* to (*μ_P_* + *μ_I_*). When the pharmacological chaperone dose is set to zero (*D* = 0), as in Masel et al. (1999), 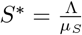. This means that when doses of pharmacological chaperones are being introduced, *S*^*^ is a different value. This *S*^*^ is less than 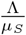 because 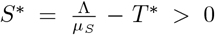. Therefore, the pharmacological chaperone treatment lowers *R*_0_ by lowering the number of susceptible PrP^C^ proteins at the PFE.

### 3.4. Endemic Equilibrium

To analyze the endemic equilibrium, assume that the interferons do not affect *R*, the non-infectious PrP^Sc^ monomers that are not joined to polymer chains (i.e. *σ* = 0). Due to the complexity of the system, the endemic equilibrium will only be analyzed in the special case where the antibody treatment is not being administered (*D* = 0).

Unlike the prion-free equilibrium and secondary infection rate, the endemic equilibrium of System 1a-1f is different from that of Masel et al. [10] when neither treatment is being administered (*D* = 0 and *μ_I_* = 0). When *D* = 0 the endemic equilibrium (EE) is given by 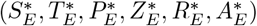 where:

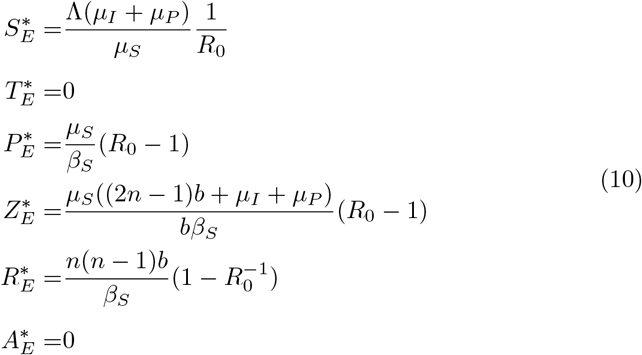

Thus the endemic equilibrium exists only if *R*_0_ > 1. Additionally, when *μ_I_* = 0, the sum 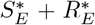 is equal to the value of *S* at the endemic equilibrium in Masel et al.’s model [10]. This occurs because PrP^Sc^ monomers are added back into the susceptible population in that model, whereas in this paper, PrP^Sc^ monomers are placed in the treatment resistant population (*R*).

### 3.5. Growth Rate

Previous models [15] have focused their analysis on the growth rate of prion concentration rather than the reproductive number *R*_0_, typical of infectious diseases. We include this analysis here for completion, as an alternative to evaluate treatment performance. The rate of exponential growth, *r*, is defined as the per capita change in number of prions chains per unit time [41]. It depends on the kinetics of the entire system and includes polymer replication and elongation. This exponential growth is the result of chains breaking into two polymers that are able to replicate. It is affected by changes in D (dosage of pharmacological chaperones) and *μ_I_* (the interferon-induced prion death rate) and is therefore useful to analyze their impact on the system.

To calculate the growth rate, it is assumed that *S, A, R*, and *T* are initially at a steady state. This reduces our original closed system of differential equations to a linear system of equations for *Z* and *P*:

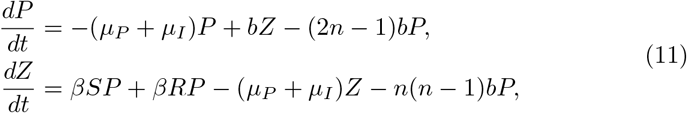

Note that the relation between *P* and *Z* is linear at all times. In fact, *Z* = *GP*, where *G* is the average chain length (see 3.3). The rate *r* then determines the exponential growth of both *P* and *Z*, and can be calculated as the dominant eigenvalue of the Jacobian matrix of System 11. The resulting equation is

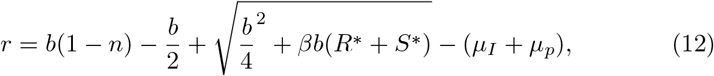

where *S*^*^ and *R*^*^ represent the *S* and *R* values of the PFE.

By replacing the value of k obtained in 4 into *r*, we obtain an expression that depends on the concentration of interferons. The exact derivation of the formula can be found in Appendix Appendix D.

As the dosage of either treatment increases the growth rate decreases. Figure 10 particularly shows how *I* and *D*, the proposed treatments, directly affect the population of prions. To examine this system, *r* is plotted against our two treatment variables: *I*, in order to study the effect of the interferon treatment, and *D*, which shows how the pharmacological chaperones affect the growth rate of PrP^Sc^, see Figure 10. It is evident that both treatments make the growth rate negative at certain concentration, however interferon treatment works better, even in low concentrations and this phenomena is related to the fact that interferons affect directly the death rate of prion proteins.

**Figure 10:**
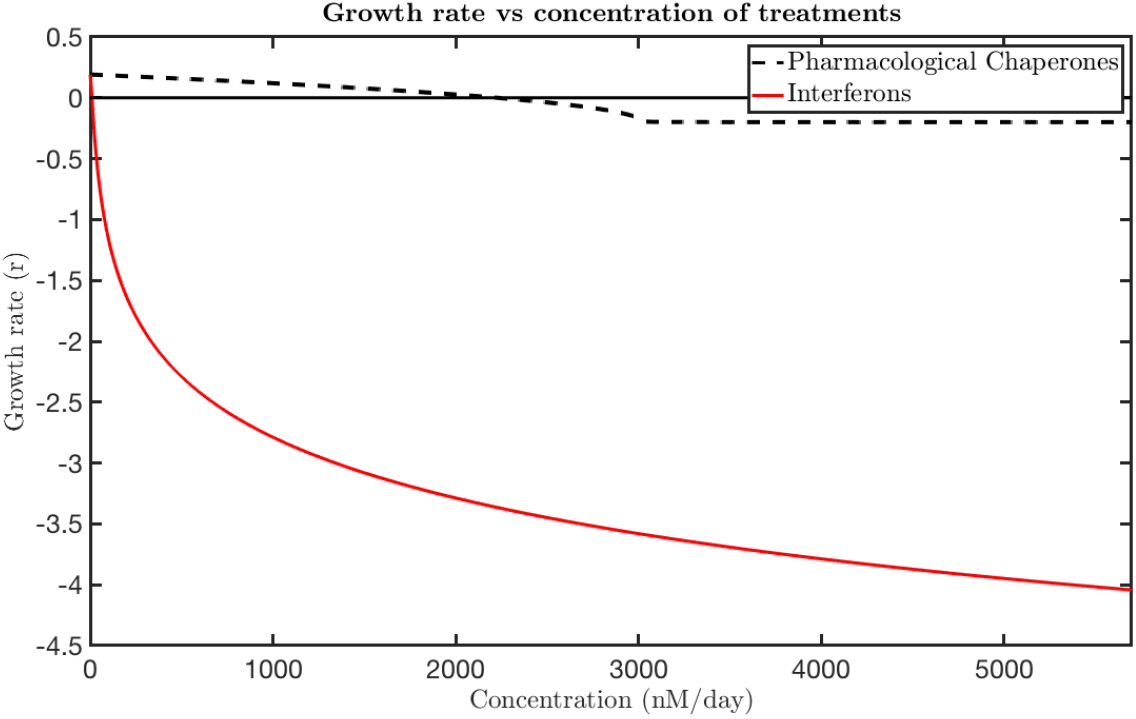
Treatment comparison based on their effect on prion growth rate. The parameters are Λ = 3000, *μ_S_* = 5, *μ_P_* = 0.2 and *n* = 2. Both treatments can achieve a negative growth rate, but interferons act at lower doses.

In Figure 11, holding *D* constant and plotting *r* against *I*, it can be seen that the growth rate decreases as *I* increases. Since the interferons remove PrP^Sc^ from the population, this limits how fast *P* can grow. Figure 11 shows three plots for three different *D*, that indicate how the system is affected with the variation of *D*. This makes biologically sense because as pharmacological chaperones are added to the system, there are fewer PrP^C^ monomers that can be added to PrP^Sc^ chains. That means that *P*, the polymers, must grow more slowly when the dosage is increased. However, the pharmacological chaperones change the growth rate much less than the interferons do, and it becomes evident that *I* reduces the growth rate much more than *D* does. This indicates that the interferon treatment reduces the prion population much more than pharmacological chaperones do. However, the dynamics of the interferon treatment affects directly the death of the prion populations (thought *μ_I_*) and the total prion death rate has the shape *μ_p_* + *μ_I_*, consequently in our model the fact that interferon can decrease the amount of prions by themselves will depend in the value of *μ_p_*. With the parameter values shown in Figure 11, then values bigger than *μ_p_* = 0.18061, interferons are able to decrease the amount of prions by themselves.

**Figure 11:**
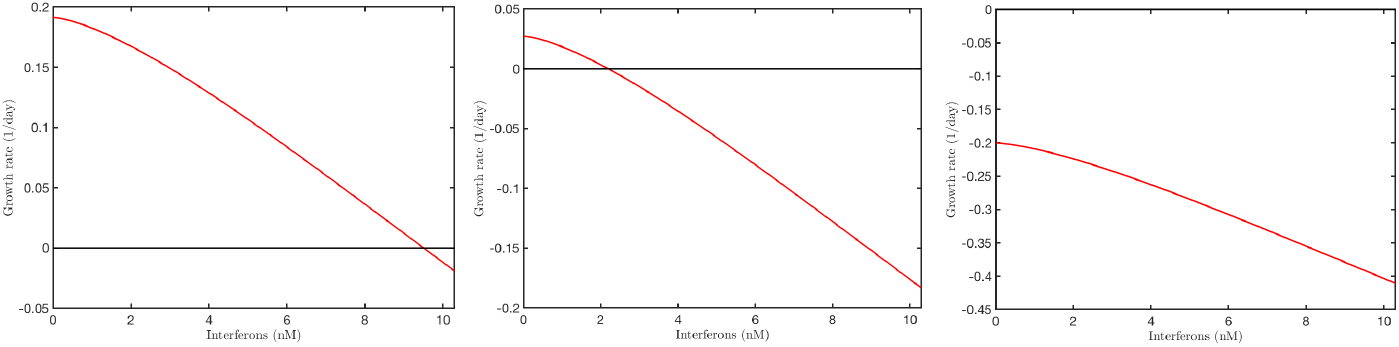
Effect of interferon dosage on the growth rate for different doses of pharmacological chaperones. From left to right: 0 nM/day, 2 000 nM/day, and 5800 nM/day. The parameters are Λ = 3000, *μ_S_* = 5, *μ_P_* = 0.2 and *n* = 2.

It was expected that the introduction of these treatments would reduce the concentration of prions in the brain. Our model shows that PrP^Sc^ decreases until it reaches the PFE by adding the treatment. From the analysis of the growth rate, we can see the first treatment has less efficacy. That is, more pharmacological chaperones are required to produce the desired result, which is to reduce the growth rate. R_0_ decreases with both treatments, increasing the pharmacological chaperones or increasing the interferons. The range for both are different; big changes in pharmacological chaperones make no significant changes in R_0_, and low changes in interferons make considerable changes in R_0_. Similar changes are evident in *r*, the growth rate of the prion population. Increasing the pharmacological chaperone dose decreases *r*, but only by a small amount. On the other hand the introduction of interferons lowers the growth rate significantly.

## 4. Numerical Results

This section includes the numerical simulations used to study the effect of treatment on prion proliferation in the brain. Parameters values are obtained from literature, particularly from Rubeinstein et al. [35]. For the parameters values see Table 3.

**Table 3:** Table of parameters for the numerical simulations. nM = nano-moles/Liter

| Name | Value | Units | Reference |
| --- | --- | --- | --- |
| Initial Conditions |  |  |  |
| $S$ | 750 | nM | [34] |
| $T$ | 0 | nM | |
| $P$ | 3 | nM | |
| $Z$ | 90 | nM | |
| $R$ | 0 | nM | |
| $A$ | 0 | nM | |
| Parameters |  |  |  |
| $\Lambda$ | 3000 | $\frac{1}{\text{day}}$ | [35] |
| $\mu_S$ | [3,5] | $\frac{1}{\text{day}}$ | [35] |
| $\mu_P$ | [0.02,0.2] | $\frac{1}{\text{day}}$ | [10] |
| $\mu_R$ | 4 | $\frac{1}{\text{day}}$ | [35] |
| $\mu_A$ | 1/29 | $\frac{1}{\text{day}}$ | [36] |
| $\mu_I$ | [0,0.2511] | $\frac{1}{\text{day}}$ | |
| $\sigma$ | [0,1] | — | |
| $\beta_S$ | 0.8 | $\frac{1}{(\text{day})\text{nM}}$ | [16] |
| $\beta_R$ | 0.8 | $\frac{1}{(\text{day})\text{nM}}$ | [16] |
| $\alpha$ | 46.656 | $\frac{1}{(\text{day})\text{nM}}$ | [37] |
| $b_{i,j}$ | 0.00032 | $\frac{1}{\text{day}}$ | [16] |
| $n$ | 2 | — | [35, 38] |
| $D$ | [0,58166] | $\frac{\text{nM}}{\text{day}}$ | [39] |

Figure 12 compares R_0_ values at time *t*_final_ with respect to different values of *D* and *μ_I_*, respectively. From this graph, it is evident that the more each treatment increases, the more *R*_0_ lowers, indicating the treatments are effective. Note that *R*_0_ decreases much more with respect to an increase in *μ_I_*, the death rate induced by interferons. This suggests that using interferons is the more effective treatment for prion diseases. It is important to note that the pharmacological chaperones affect R_0_ as well. However, these results indicate that the secondary infection rate is much more sensitive to the changes in *μ_I_*, a sign that treatment of prion diseases with interferons may be more useful.

**Figure 12:**
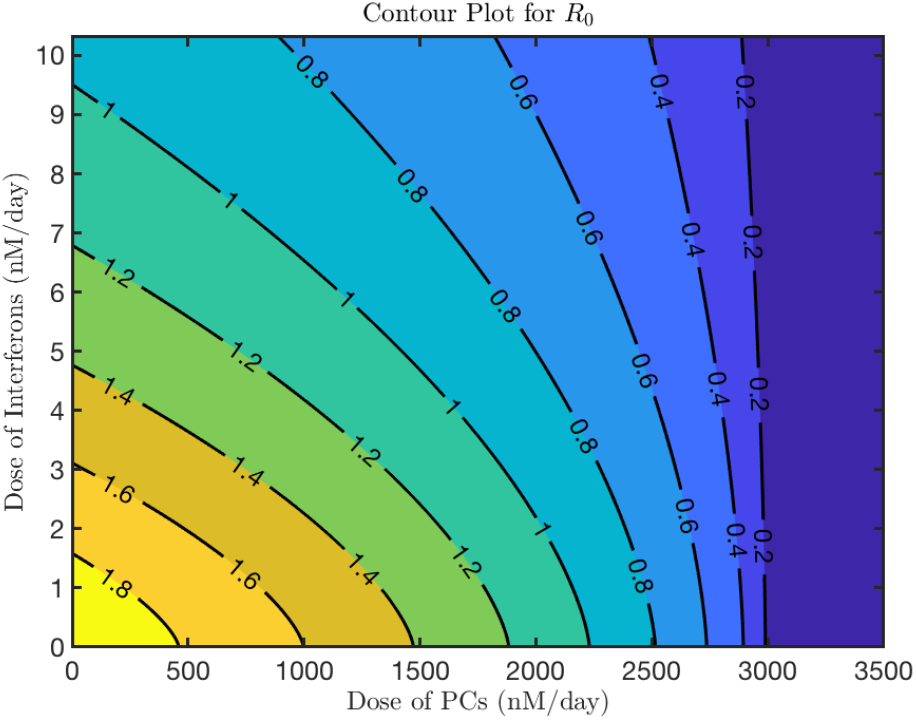
Contour plot of R_0_ vs. Interferons (*μ_I_*) and Pharmacological Chaperone Dosage (*D*). The parameters are Λ = 3000, *μ_S_* = 5, *μ_P_* = 0.2 and *n* = 2. The graph after 3100 of Dosage keeps constant.

Figure 13 shows the relationship between the endemic value of *P* compared to the treatment levels is evident. When no treatment is applied, *P_E_* is at its highest in the lower left corner of the figure. This makes sense; treatments should limit the strength of the final infection level. Travelling along the *I* axis, it is observed that as *P_E_* decreases more steeply the interferon treatment increases. This makes sense with the previous analysis of *r*, the growth rate; the interferon treatment is the more effective of the two treatments. The decrease that happens along the *D*-axis is much less pronounced, but still existent. Note that the pharmacological chaperone dosage is limited due to toxicity of the treatment. Pharmacological chaperones affect the amount of PrP^Sc^ as well, though not as much the interferons do.

**Figure 13:**
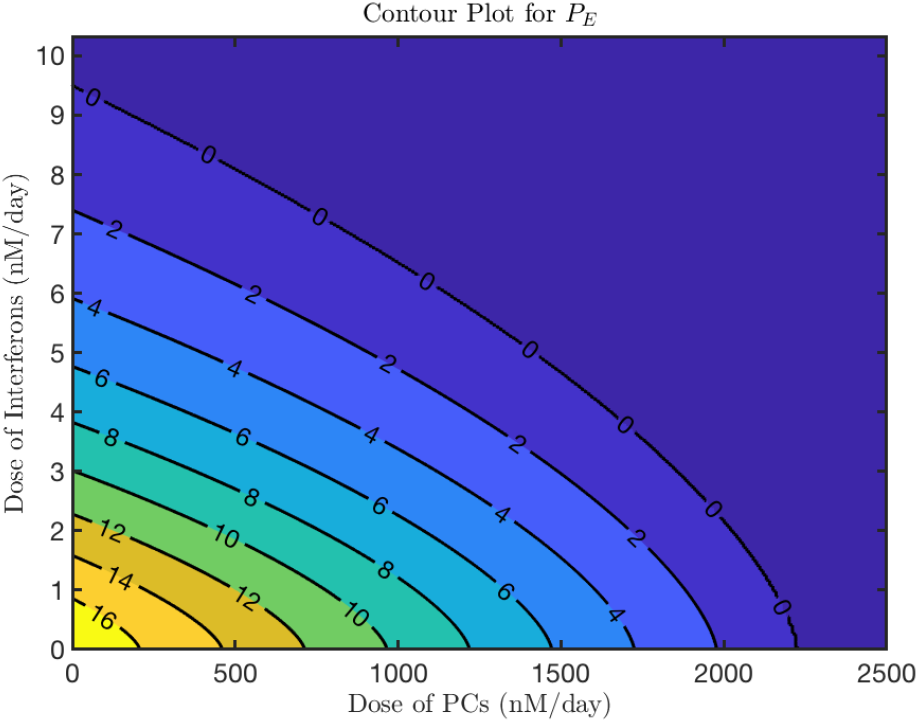
Contour plot of *P_E_* vs. Interferons (*I*) and Dosage (*D*). The parameters are Λ = 3000, *μ_S_* = 5, *μ_P_* = 0.2 and *n* = 2.

Figure 14 effectively shows the synergistic effect of the two treatment’s on the endemic equilibrium (*P_E_*). While interferons are more effective than pharmacological chaperones at both reducing the value of the endemic equilibrium and the time it takes to reach that equilibrium, the combined treatment shows a near 100-fold decrease in endemic equilibrium than that of the untreated infection.

**Figure 14:**
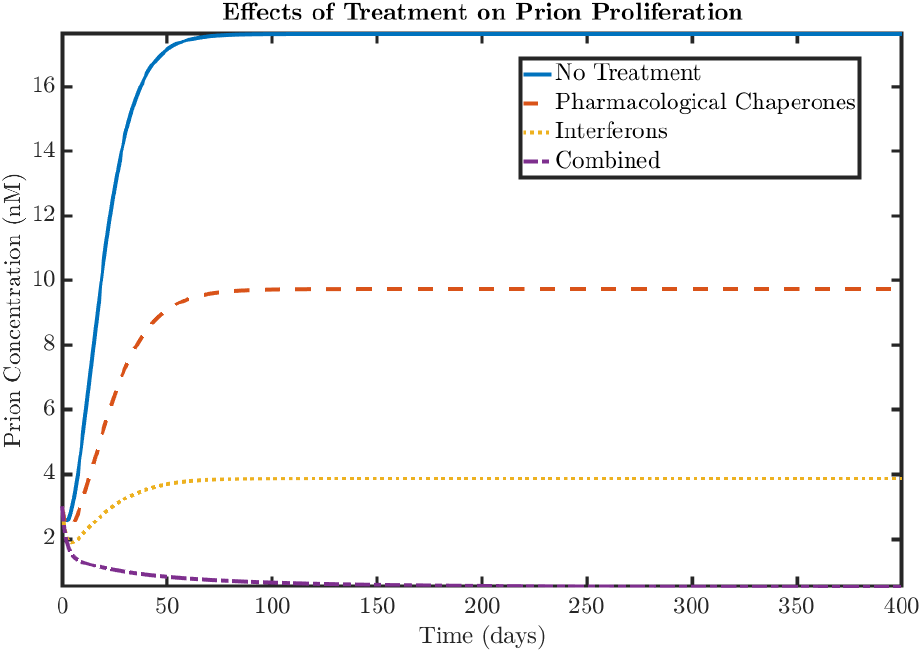
This figure shows a special case of Figure 12. Prion concentration changes over time with various treatments implemented. Pharmacological chaperones and interferons are represented at, 1000 nM/day, and 6 nM/day respectively. Each of these dosages alone will go to endemic equilibrium. This figure shows that the treatments are able to be combined at the same dosages to go to disease free equilibrium. With no treatment, the prion concentration increase faster than with treatments. (For parameters, see Table 3). The parameters are Λ = 3000, *μ_S_* = 5, *μ_R_* = 5, *μ_P_* = 0.2 and *n* = 2.

Figure 15 shows the differing concentration of prions at a range of constant interferon dosages. It can be shown that at these particular initial conditions found in Table 3, there is a transition of stable equilibrium from a stable endemic equilibrium at low dosages of interferons to a stable prion-free equilibrium at higher dosages. As the interferon dose increases, the prion concentration does not increase as quickly as without treatment. The latter two graphs in the figure shows how the final concentration of prions depend on interferons; this value goes to zero as we increase both interferon dose and, relatedly, the interferon-induced death rate. This treatment, at high enough doses, appears to work against prion proliferation.

**Figure 15:**
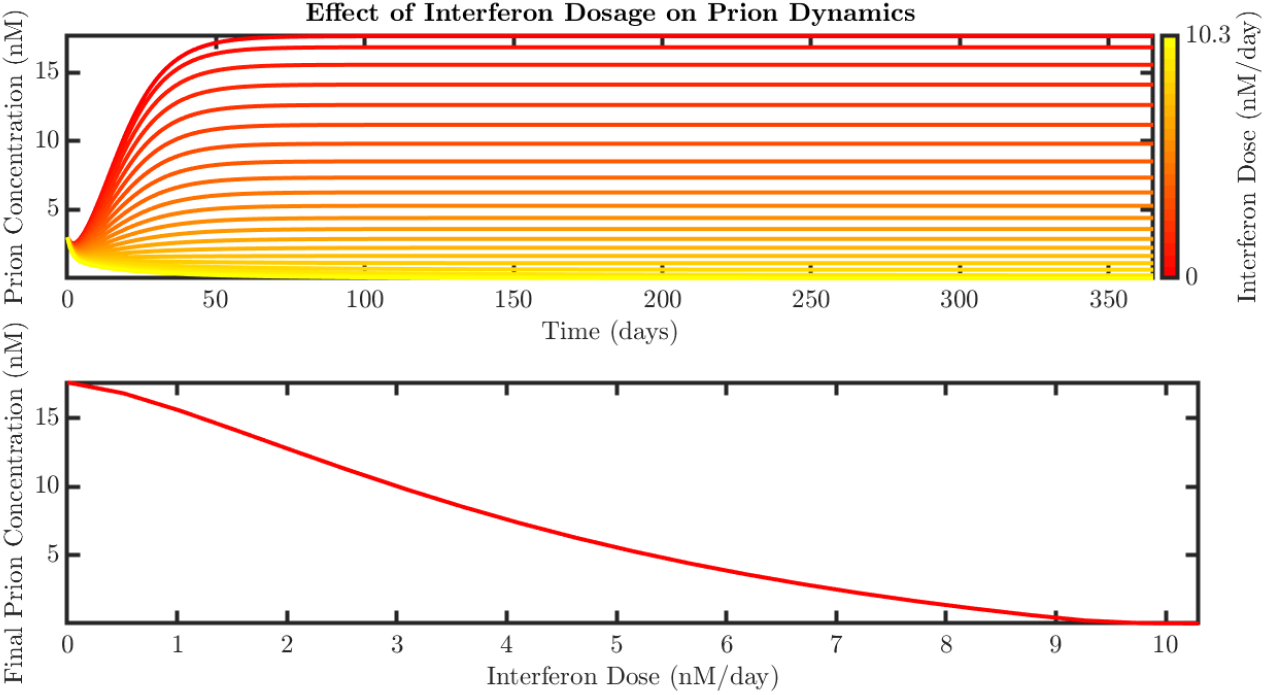
**Top:** The concentration of prions as a function of time, where each line is a different dose of Interferons. **Bottom:** The final prion concentration as a function of dose of Interferons (For parameter values, see Table 3). The parameters are Λ = 3000, *μ_S_* = 5, *μ_R_* = 5, *μ_P_* = 0.2, *n* = 2 and *D* = 0.

The prion concentration over time with different constant dosages of pharmacological chaperones and no interferon treatment was also examined. Figure 14 shows how combination of lower-dose treatments can be effective. In this graph, the administered dosages of both pharmacological chaperones and interferons alone cannot bring the prion population to zero. Each lowers the endemic equilibrium but does not eliminate the prion population. However, if the low doses of both treatments are combined, they can work together to bring the prion concentration down together. The synergistic effect is significant.

**Figure 16:**
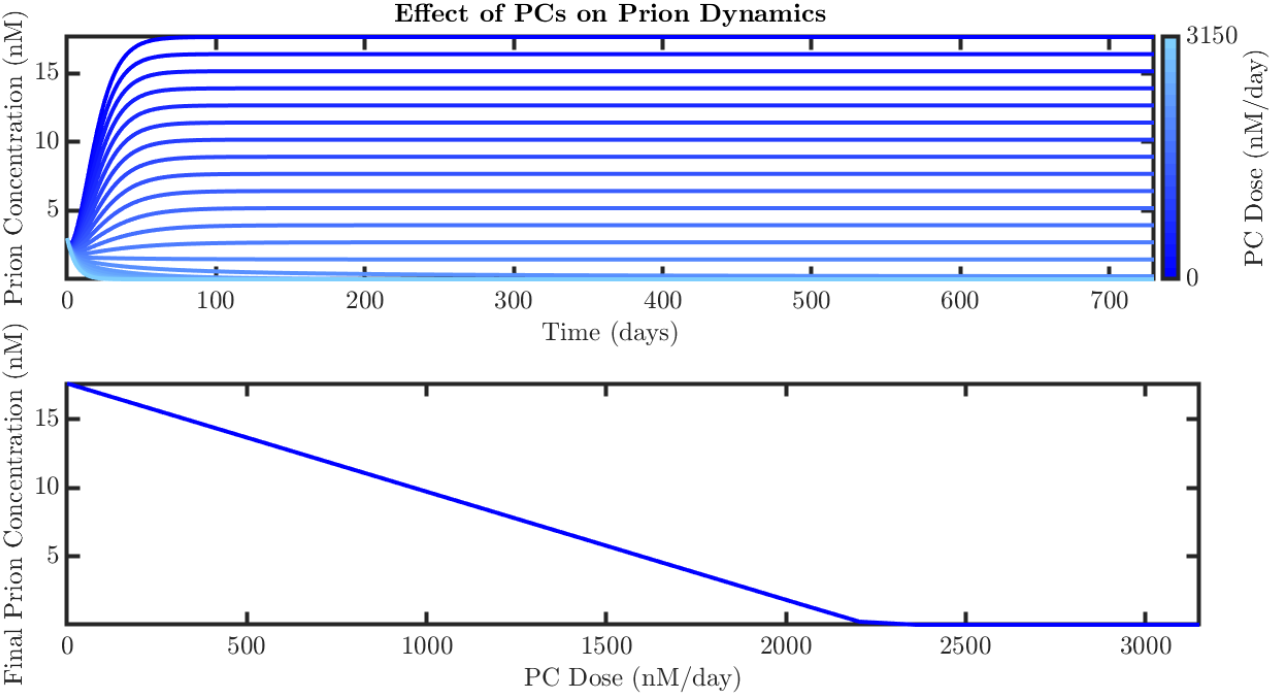
**Top:** The concentration of prions as a function of time, where each line is a different dose of pharmacological chaperones. **Bottom:** The final prion concentration as a function of dose of pharmacological chaperones. (For parameter values, see Table 3). The parameters are Λ = 3000, *μ_S_* = 5, *μ_R_* = 5, *μ_P_* = 0.2, *n* = 2 and *μ_I_* = 0.

Figure 17 shows the combination of interferon and pharmacological chaperone treatments. The red and orange lines are the same as in Figure 15. The superimposed blue and purple lines show the effect of introducing a constant dose of pharmacological chaperones while varying interferon dose. The upper limit for the prion concentration is much lower than with a single treatment. The dual treatment also slows the growth of PrP^Sc^ more than interferons alone does. The second and third graphs in Figure 17 show how interferons affect the final prion concentration, again with both no pharmacological chaperones (red) and a constant dose of pharmacological chaperone treatment (blue). The addition of a constant treatment dose means that the final prion concentration is achieved sooner. This graph shows us that using the two treatments is a way to reduce prion concentration more and faster than either pharmacological chaperones or interferons alone.

**Figure 17:**
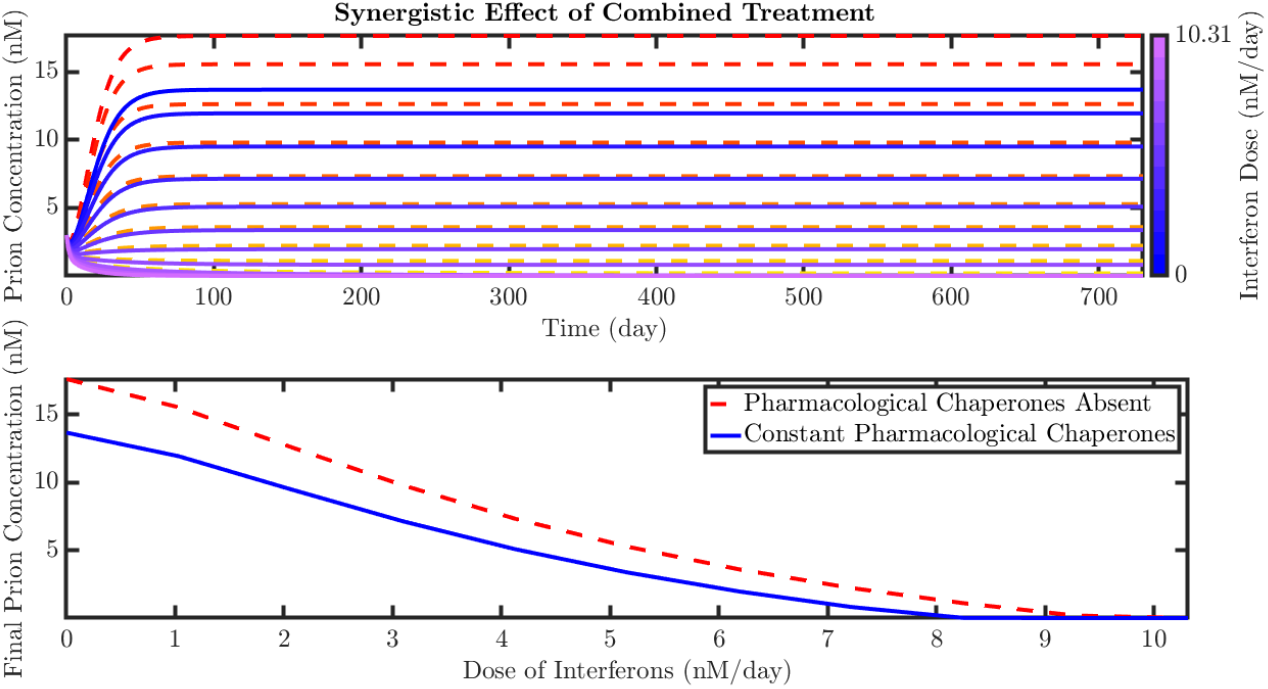
Combined Interferon and Pharmacological Chaperones. **Top:** The concentration of prions as a function of interferon and pharmacological chaperone doses. Red is only pharmacological chaperone doses, while blue represents varying interferon doses and constant pharmacological chaperones. Pharmacological chaperons are at a constant dose of 500 nM, which is right at the toxic threshold. **Bottom:** The final prion concentration as a function of dose of interferons, with and without the same doses of pharmacological chaperones as above. The parameters are Λ = 3000, *μ_S_* = 5, *μ_R_* = 5, *μ_P_* = 0.2 and *n* = 2.

## 5. Discussion

In this paper, previous models by Nowak et al. [12] and Masel et al. [10] on prion dynamics in the brain were examined. Our work introduces two possible treatments for prion diseases. The first treatment uses pharmacological chaperones, which prevent PrP from misfolding into PrP^Sc^. The other treatment uses interferons, a signalling part of the immune system that reduces the PrP^Sc^ population. This work examined how these treatments affected the population of prions in the brain. From the results of the numeric simulations it was clear that both treatments have the potential to reduce prion proliferation. However, through analysis of the basic reproduction number (R_0_) and the prion growth rate (*r*), it can be seen how the treatments work in tandem.

Compared to the pharmacological chaperones, the interferons bring the prion growth rate to negative values at lower doses and achieve a faster exponential decay at high doses. The pharmacological chaperones, although capable of inducing exponential decay in the PrP^Sc^ population, have a limited capacity to increase the magnitude of the decay rate. However, beneficial effects were evident when studying the model with combined treatments. The addition of pharmacological chaperones to the interferon treatment reduces the interferon dose required to bring the prion growth rate to negative values. As a consequence, the combination of treatment doses that would independently be insufficient to prevent prion accumulation can effectively induce an exponential decay to undetectable values.

Further experimental work is required to determine the best pharmacological chaperone to use in a monomer-blocking treatment, as well as the most convenient interferon type for human administration. The range of viable dosages for humans may vary significantly according to the agent used. Therefore, rather than recommending a treatment with the specific pharmacological chaperone and interferon used as reference here, we provide a quantitative framework to analyze the interaction between the two treatments while taking into account the toxic dose of each drug. Toxic thresholds can be readily adjusted in our model to match experimental conditions and determine the medically feasible dosage ranges for the system to evolve.

It is important to note that the work presented here does not intend to provide a cure for prion diseases, but rather to guide future experimentation through treatment simulation and analysis. Interferons and pharmacological chaperones have not been shown to eliminate the disease altogether. Further research beyond this work is needed if any steps are to be made towards a cure. Mathematically, the optimization of the two treatments would be useful. Looking for specific dosages of both pharmacological chaperones and interferons and best timing of these doses is also important. The results in this paper would be best supported by a toxicity optimization; like all drugs, the treatments presented here can not be given to a patient freely. What would be the best combination of treatment to reduce harm done by the drugs while still affecting the prion population? Additionally, further research of other possible treatments and treatment combinations is essential. Pharmacological chaperones and interferons are not the only two possible ways to treat prion diseases, and it is important to take into account all possible therapies. *In vivo* experiments are necessary to show if these treatments actually have any affect on prion diseases. Once more substantial research has been done, a cost analysis of these treatments would also be useful to reduce cost for the patient. Prion diseases are still fatal, still dangerous, still incurable. But research is happening, and perhaps we are one step closer to solving the mystery of these strange diseases.

## 6. Acknowledgments

We would like to thank Dr. Carlos Castillo-Chavez, Founding and Co-Director of MTBI, for giving us the opportunity to participate in this research program. We would also like to thank Co-Director Dr. Anuj Mubayi, as well as Coordinators Ms. Rebecca Perlin and Ms. Sabrina Avila. We also want to give special thanks to Dr. Susan Holechek. This research was conducted as part of 2018 MTBI at the Simon A. Levin Mathematical, Computational and Modeling Sciences Center (MCMSC) at Arizona State University (ASU). This project has been partially supported by grants from the National Science Foundation (NSF - Grant MPS-DMS-1263374 and NSF – Grant DMS-1757968), the National Security Agency (NSA – Grant H98230-J8-1-0005), the Office of the President of ASU, and the Office of the Provost of ASU.

## Appendix A. Mathematical Proofs for the Equations

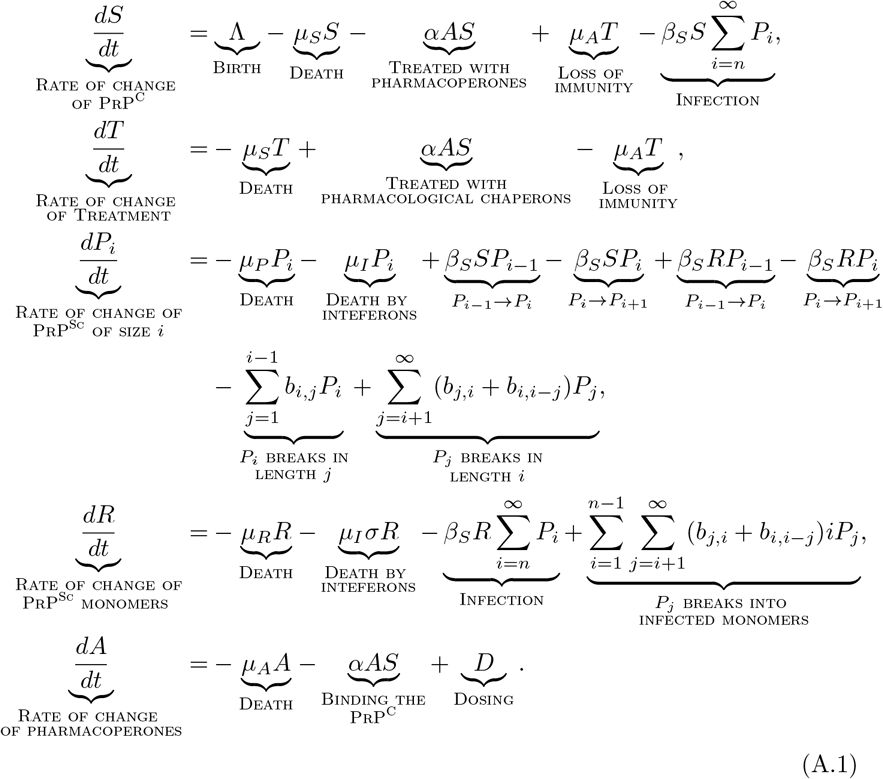

### Appendix A.1. Derivation of the Infinite System of Differential Equations

In this section, the derivation of the system of differential equations from the infinite dimensional system is explained. First, take the term

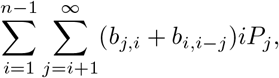

which describes a polymer of length *i* splitting into two polymers, with one of them of length less than *n*. When the shorter polymer is below the *n* threshold, it dissociates into infected monomers, and so the monomers go from the *P_i_* class to *R*. Thus, for a single length *j* we can represent the monomers flowing into *R* by the following:

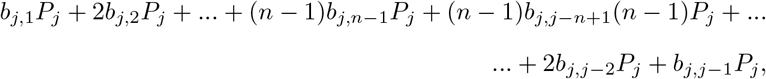

The sum of this value over all lengths *j* can be written,

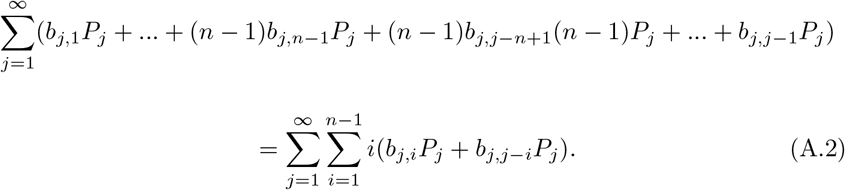

It is known that *b_j,i_* = 0 if *i* ≥ *j*; so Equation A.2 can be rewritten as:

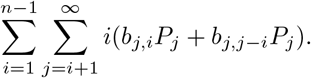

Different assumptions about the dynamics of aggregate growth and fragmentation can be embodied in the matrix *b_i,j_*. In this, we assume that *b_i,j_* = *b*, which changes the sum as follows:

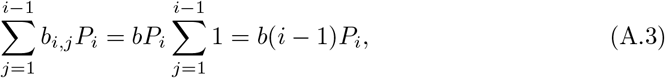

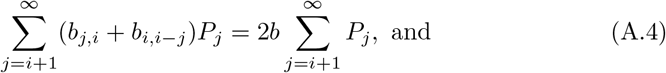

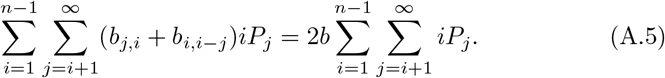

Therefore, the construction of the infinite dimensional system can be written as:

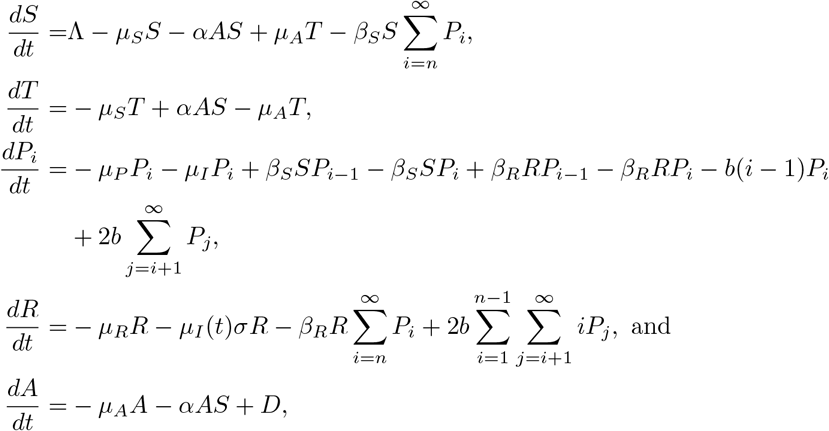

### Appendix A.2. Closing the System of Differential Equations

Here we close the system with infinite equations by summing over *P_i_*. Let

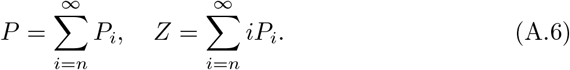

Therefore, it is assumed that these sums are convergent since in any biological system, there will only be a finite number of prions. Their derivatives can be written as 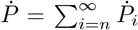 and 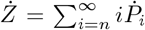. The derivative of *P* can then be reduced as follows:

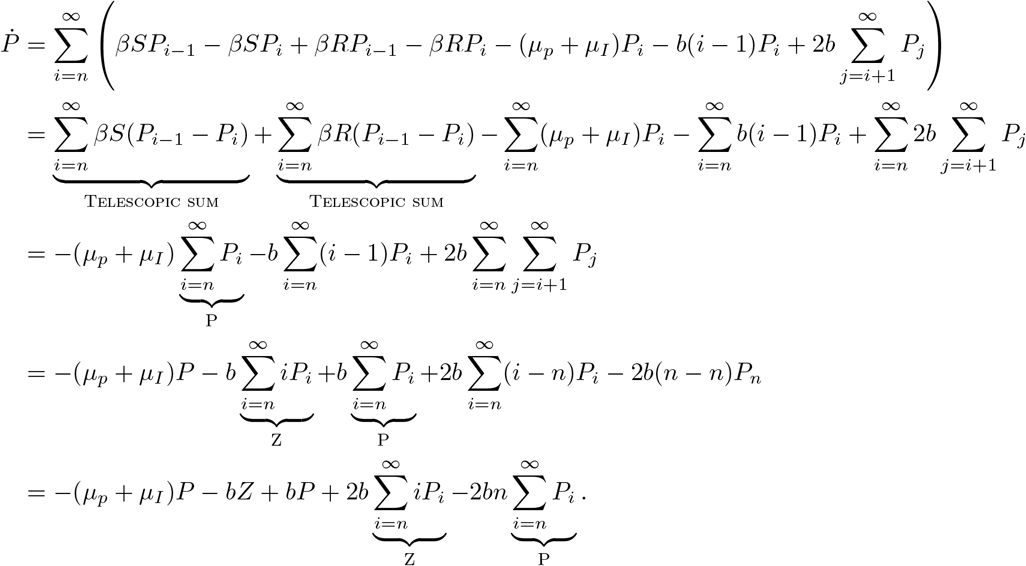

Thus,

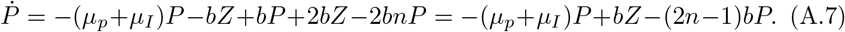

The derivative of *Z* can also be rewritten:

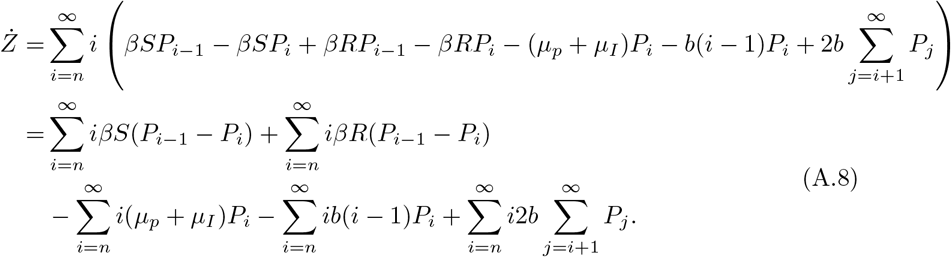

The sums in Equation A.8 can be reduced as follows:

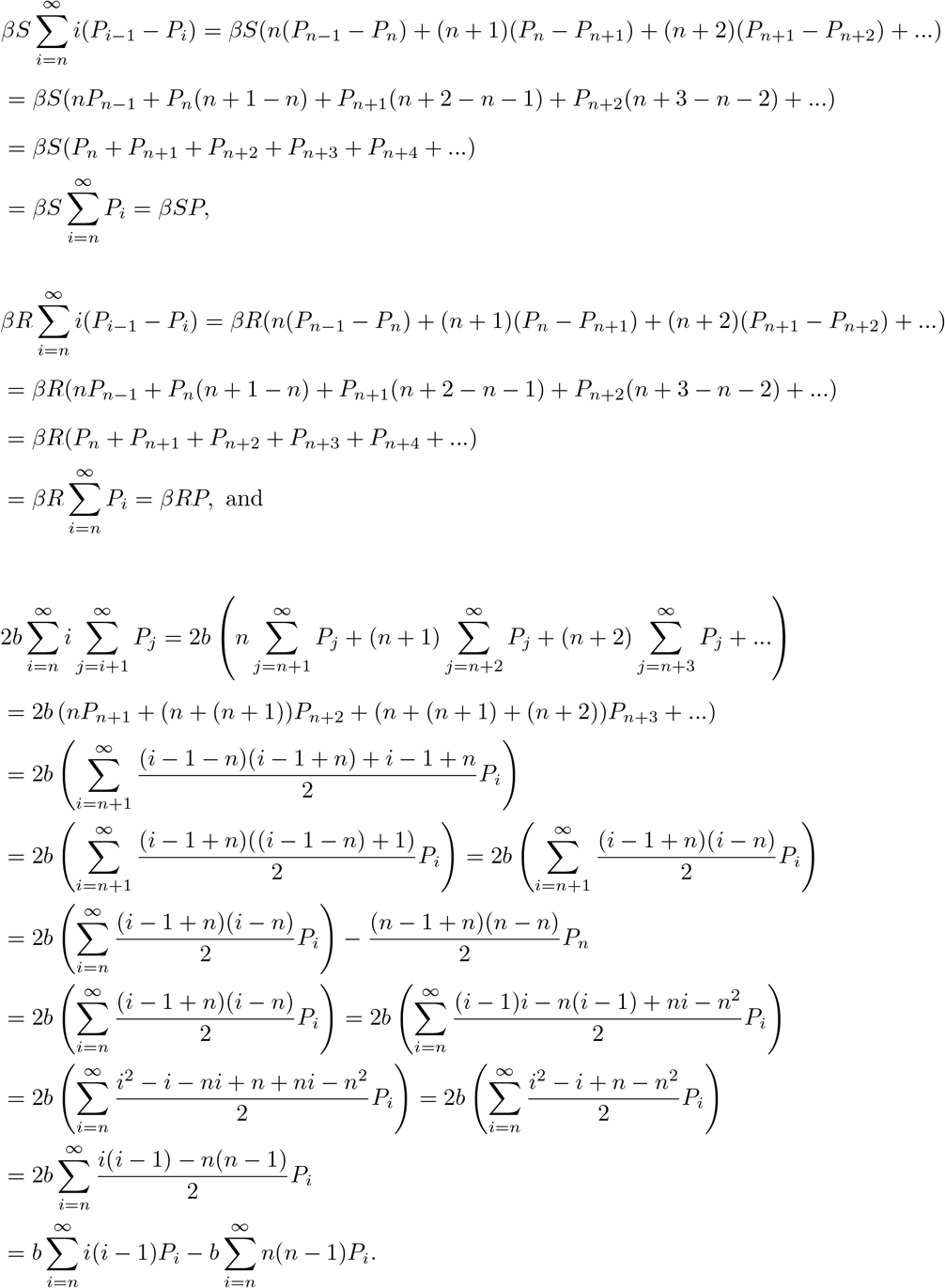

Plugging these reduced terms back into *Ż* yields

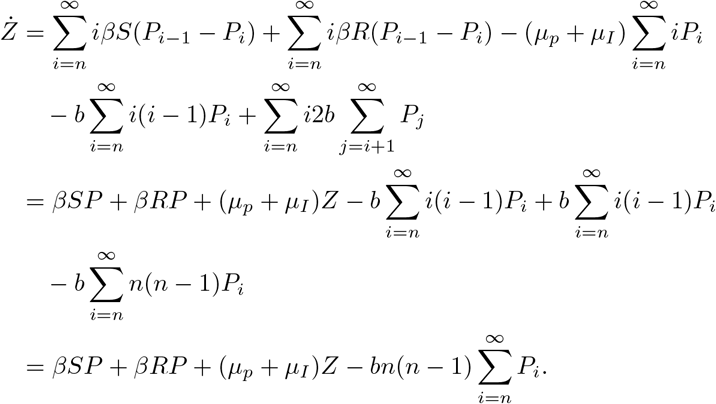

Thus,

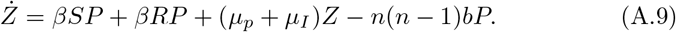

Now the infinite sums in Equation 2.2e (shown here in Equation A.10) can be reduced.

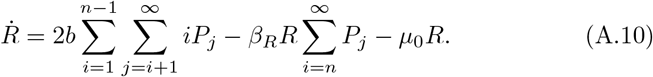

These sums can be reduced as follows:

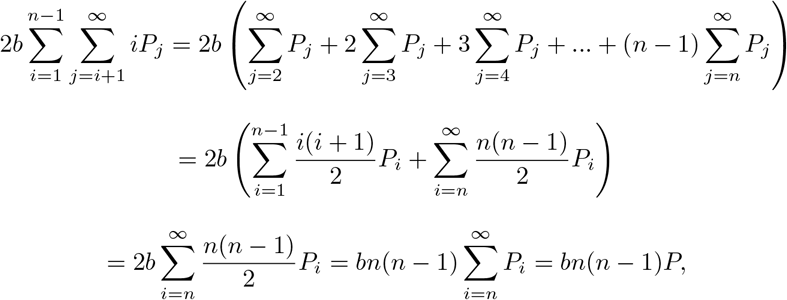

because *P_i_* = 0 for *i* < n. Thus,

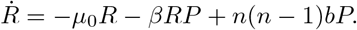

### Appendix A.3. Stability Condition of the Prion Free Equilibrium

The linearization matrix **J**(**E**^*^) of System 2.2 around *E*^*^ = (*S*^*^, *T*^*^, 0,0,0, *A*^*^) is

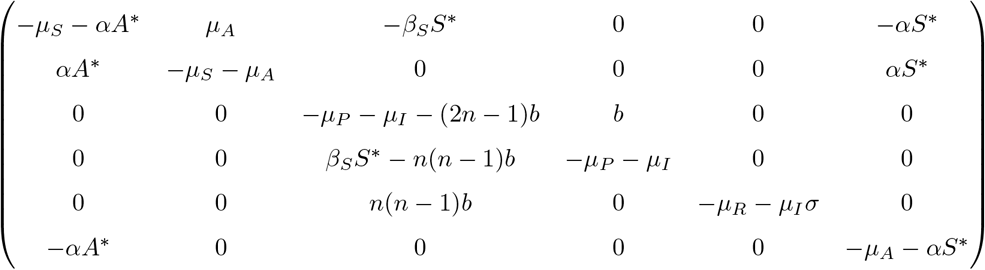

The eigenvalues of the matrix *J*(*E*^*^) are:

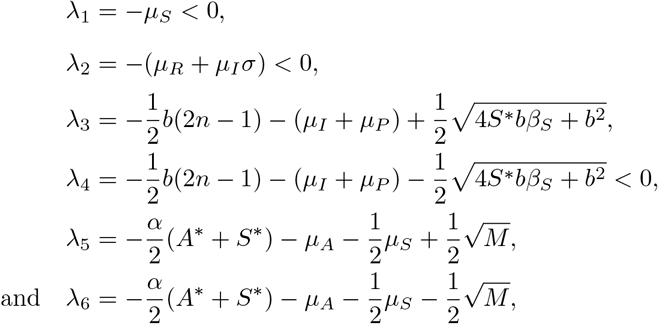

where

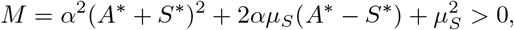

given that

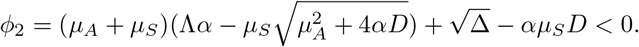

i. From the value of λ_3_, we have the λ_3_ < 0 when *ϕ*_4_ < 0, where

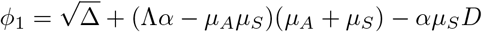
ii. From the value of λ_4_, we have the λ_4_ < 0 if 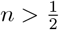.
iii. From the value of λ_5_, because 4*μ_A_* (*μ_S_*+*μ_A_*)+4*α*(*μ_S_S*^*^ + *μ_A_S*^*^ + *μ_A_A*^*^) > 0, so λ_5_ < 0. But, we want *M* is positive. Hence, we need *ϕ*_2_ < 0.

As a result, equilibrium *E*^*^ is locally stable when *ϕ*_1_ < 0(*R*_0_ < 1).

Proof:(i) if λ_3_ < 0,we can obtain 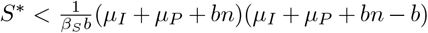, yield

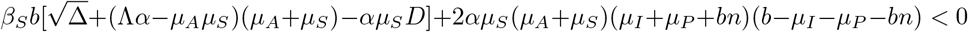

we can let 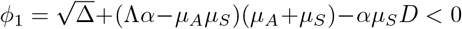 and *b*(*n*–1)+*μ_I_*+*μ_P_* > 0, then λ_3_ have negative real part.

(iii) Because 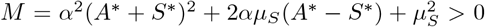. We just need 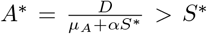, then *M* is positive. So, *S*\* satisfy *αS*^2^ + *μ_A_S* – *D* < 0, because 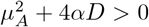 and –*D* < 0, hence, this quadratic equation must have positive root 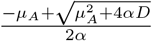. Let 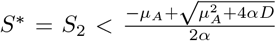. We can obtain *ϕ*_2_ < 0.

### Appendix A.f. R_0_ Using Next Generation Matrix

This section will show in detail how 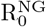 was calculated using a Next Generation Matrix. System 2.2 is rewritten as 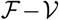 where the terms in 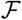 represent the creation of new infectious prion chains. Therefore,

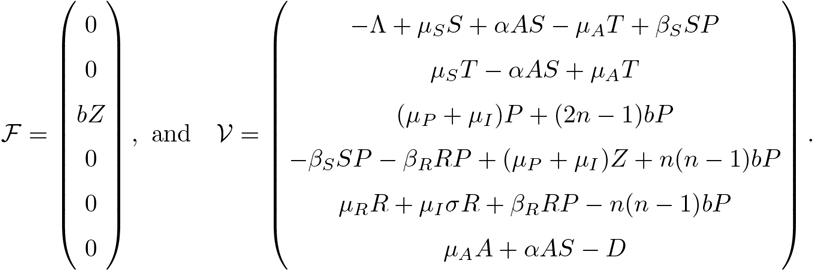

Next, take the jacobian of 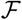 and 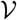 evaluated at the prion-free disease equilibrium (*S*^*^, *T*^*^, 0, 0, 0, *A*^*^):

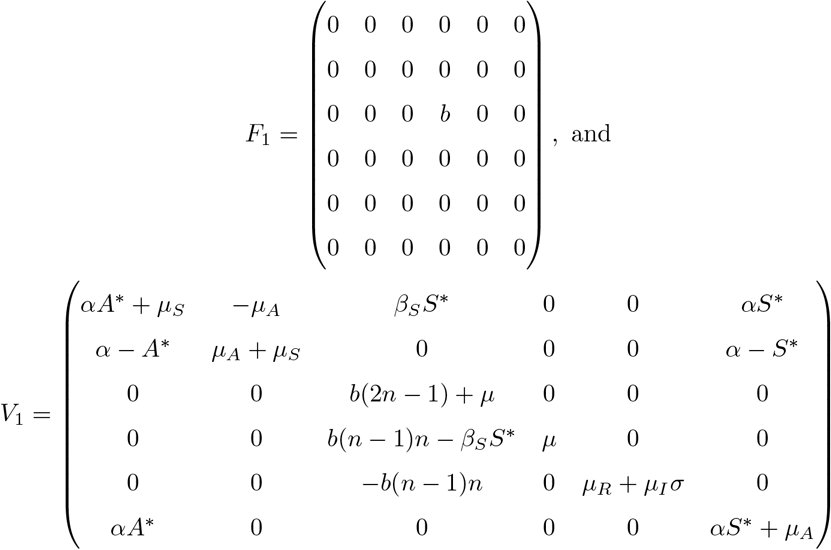

where *μ* = *μ_I_* + *μ_P_*. Now, *F* and *V* can be redefined to reduce the system:

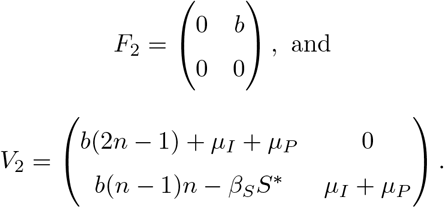

The inverse of *V*_2_ can be written as:

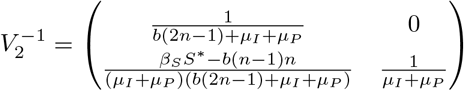

Now, find the value 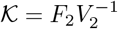:

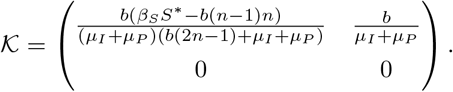

Therefore, 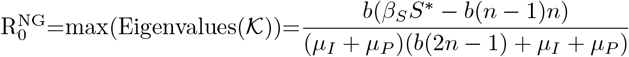.

## Appendix B. Derivation of Interferons Formula

### Appendix B.1. Type I IFN-B inhibit prion propagation in infectious cells and animal models

The values of the concentrations of interferon’s were taken from experimental data [31]. In the experiments,the concentration is measured in *kU/ml* and 48h after interferon doses the rate is considered dividing the total amount of prion over the control amount (which does not include the interferon treatment).

Pairs of experimental values:

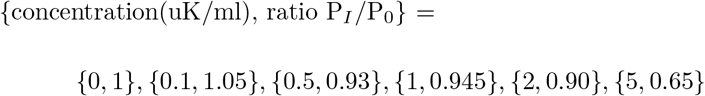

#### Appendix B.1.1. Unit Conversion

Since the experimental data is in **kU/ml** we need to do a conversion of units to obtain the units **nM/L**.

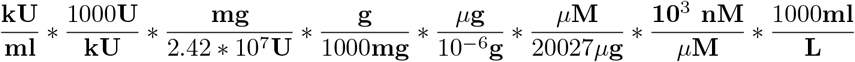

where, 20027**g/mol** is the molecular mass of an interferon [42]. Then the new list of pairs of values {concentration(nM/L), ratio*P_I_*/*P*_0_} is

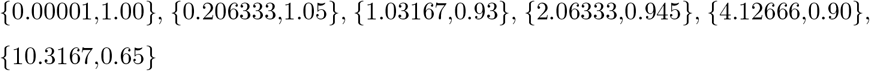

#### Appendix B.1.2. Interpolation

The interpolation of the data with a Hill Equation is described by the equation

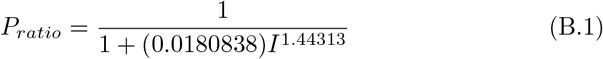

Then, assuming that the dosage *D* and the death rate caused by interferons *μ_I_* is constant, then we rewrite the total dead rate of the polymer chain *μ_P_* ; *μ_I_* as (1 + *k*)*μ_P_*. We assume that the behavior of prions with respect to time (including the treatment) is given by an exponential decay:

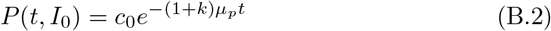

Then, the ratio between the amount of prions when there is treatment and when there is not treatment is given by

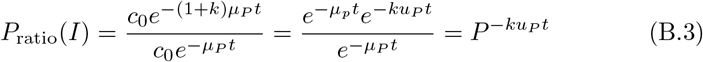

Then, if it is assumed a *t* = 2 days (48 hours) and it is solved for *k*, it is obtained:

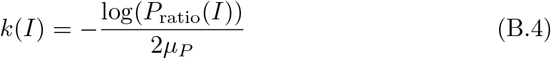

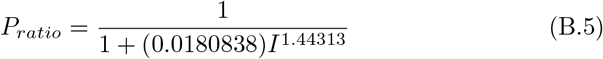

Where *P*_ratio_(*I*) will be a ratio between 0 and 1.

##### Code

The code for this calculations was created with Wolfram Mathematica.

~~~
(*Values taken from the experiment*)
rateI4={{0.001,1.00},{0.206333,1.05},{1.03167,.93},{2.06333,.945},
{4.12666,.90},{10.3167,.65}};
CvsI= ListPlot[ratel4]
~~~

~~~
hillmodelinter=(1/(1+(x/a)^n));
hillfit=FindFit[ratel4, {hillmodelinter,{n>0,a>0}},{a,n}, x]
Show[Plot[hillmodelinter/.hillfit, {x, 0, 9.5*10^6}],
ListPlot[rateI4, PlotRange -> Automatic]]
~~~

~~~
(*Assumptions*)
p[t_,co_,k_,uo_]:= co E^(-(1+k)uo t)
po[t_,co_,k_,uo_]:= co E^(-uo t)
inter[Io_,r_,t_]:= Io E^(-r t)
(*k[Io];*)
(*Assume ut and dt ctes?*)
~~~

~~~
(*Assuming a t=2*)
ratio2= FullSimplify[(p[2,co,k,uo])/(po[2,co,k,uo])]
~~~

~~~
(*Assuming a t is a variable*)
ratio=FullSimplify[(p[t,co,k,uo])/(po[t,co,k,uo])]
k2=Assuming[
  { { k, uo,f2} \[Element] Reals, f2>0},
  FullSimplify[Flatten[{
   Solve[ratio2==f2,k,Reals]},2]
  ]
]
kt=Assuming[
  { { k, uo} \[Element] Reals, f2>0,f2<1 },
  FullSimplify[
   Solve[ratio2==f2,k,Reals]
  ]
]
kIt=Assuming[
  { { k, uo} \[Element] Reals, f>0,f<1, E^(r t)>0, Io>0},
  FullSimplify[
   Solve[ratio2==inter[Io,r,t],k,Reals]
  ]]
pratio=hillmodelinter/.hillfit
~~~

## Appendix C. Interferon Analysis

There are several experimental treatments that aim to stop prion proliferation, one of which is the direct dosing of interferons. Experiments done by Ishibashi et al. (2019) [31] have indicated that the interferon signalling interferes with prion propagation. In this research, a group of mice was infected with prions and then treated with different concentrations of interferons. The concentrations of prions was measured after 48 hours of the inoculation of interferons. Due to the interferon signalling, there was a decrease in the prion population.

From this experiment, the following experimental data was obtained:

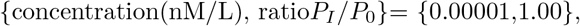

For each of the samples treated with different concentrations, it is measured the ratio *P_ratio_* = *P_I_*/*P*_0_ between the amount of prions before and after the interferons were introduced. Then, the data was interpolated with a Hill equation as shown in Figure C.18.

**Figure C.18:**
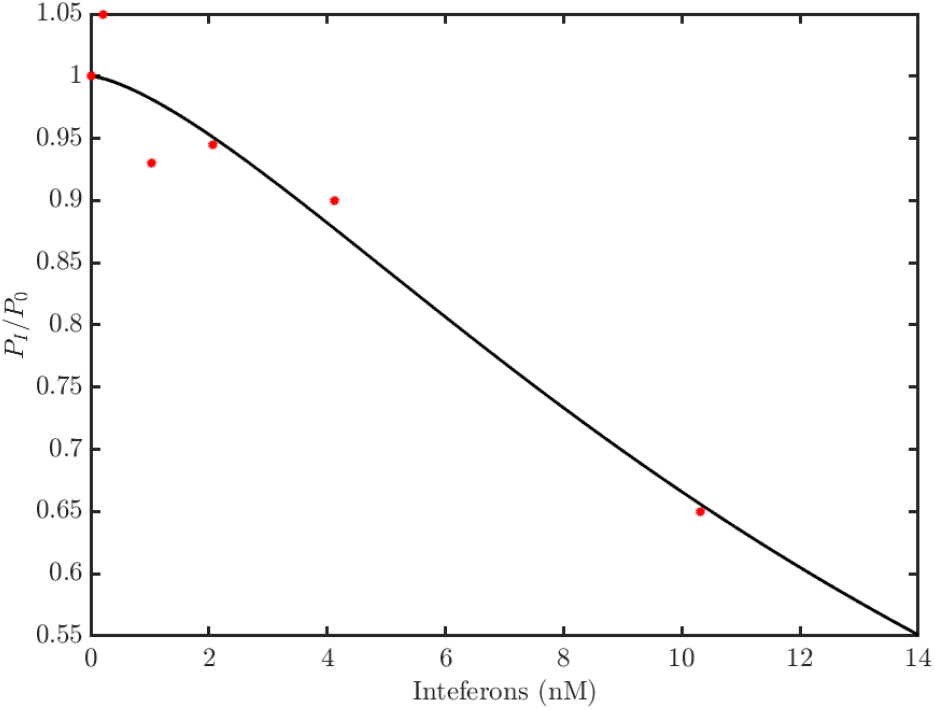
*P_I_*/*P*_0_ vs concentration. The ratio of prions measured in experiments with respect to the different concentrations of interferons. A Hill equation was used to interpolate the data. The range of the interferons is [0,10.31]

From this interpolation the following equation was obtained.

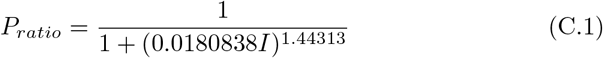

where *I* is the concentration of interferons. Then, assuming that the dosage *D* and the death rate caused by interferons *μ_I_* are constant, we rewrite the total death rate of the polymer chain *μ_P_* + *μ_I_* as *μ_P_* + (*k*)*μ_P_*. We assume that the behavior of prions with respect to time (including the treatment) is given by an exponential decay:

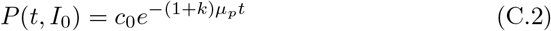

The ratio between the amount of prions when there is treatment and when there is not treatment is given by

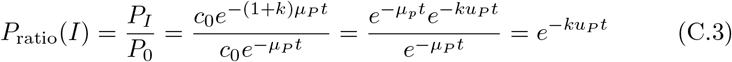

Because the experiment dosed interferons over 48 hours, we assume *t* = 2 days and solve for *k*.

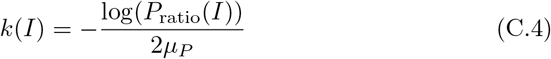

where *P*_ratio_(*I*) is a ratio between 0 and 1 defined by equation C.3.

Note that the experimental data *I* was a unique dose. The *P_ratio_* values were measured after two days of inoculation. It is assumed for calculations that the dose per day is the half of the total dose.

## Appendix D. Growth Rate

### Appendix D.1. Getting r

In this section of the appendix we show how to get the growth rate. Asume that *S, T, R* and *A* are initially in a steady state. The system if *P* and *Z* is now linear, and will either accumulate or decay exponentially at exponentially at the rate according to the dominant eigenvalue of the Jacobian matrix. After the average size reaches equilibrium, exponential growth in the abundance of PrPSc over time t occurs according to

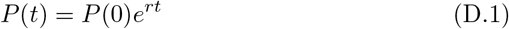

and

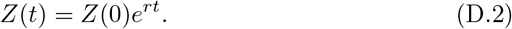

Taking the equations for *P* and *Z*,

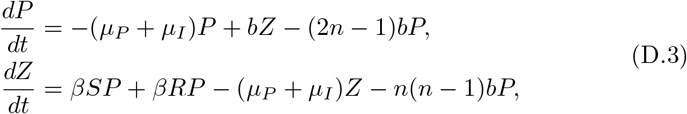

getting the Jacobian that is given by

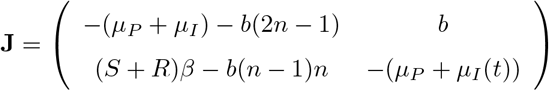

The characteristic polynomials of **J** that is given by

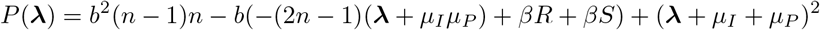

and the solutions are given by

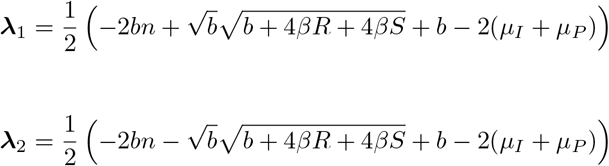

taking maximum eigenvalue

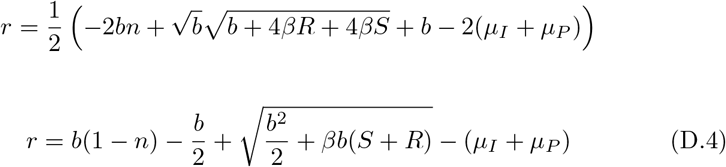

Then, the values of the Prion Disease Equilibria for *S*\* and *R*\* are introduced from equations 7. Then, the following expression is obtained:

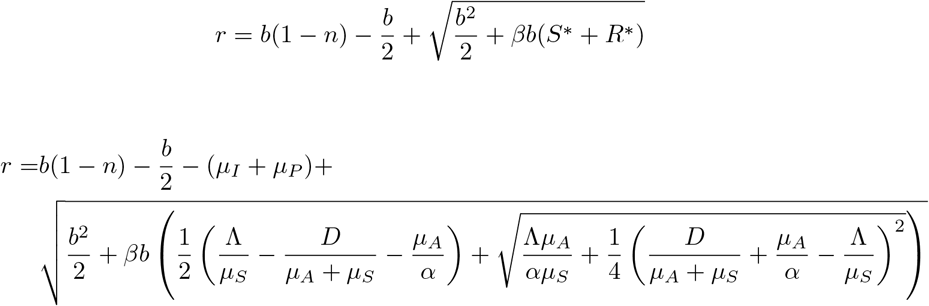

An illustration of the growth rate as a function of the dose of pharmacological chaperones D can be found in Figure D.19

**Figure D.19:**
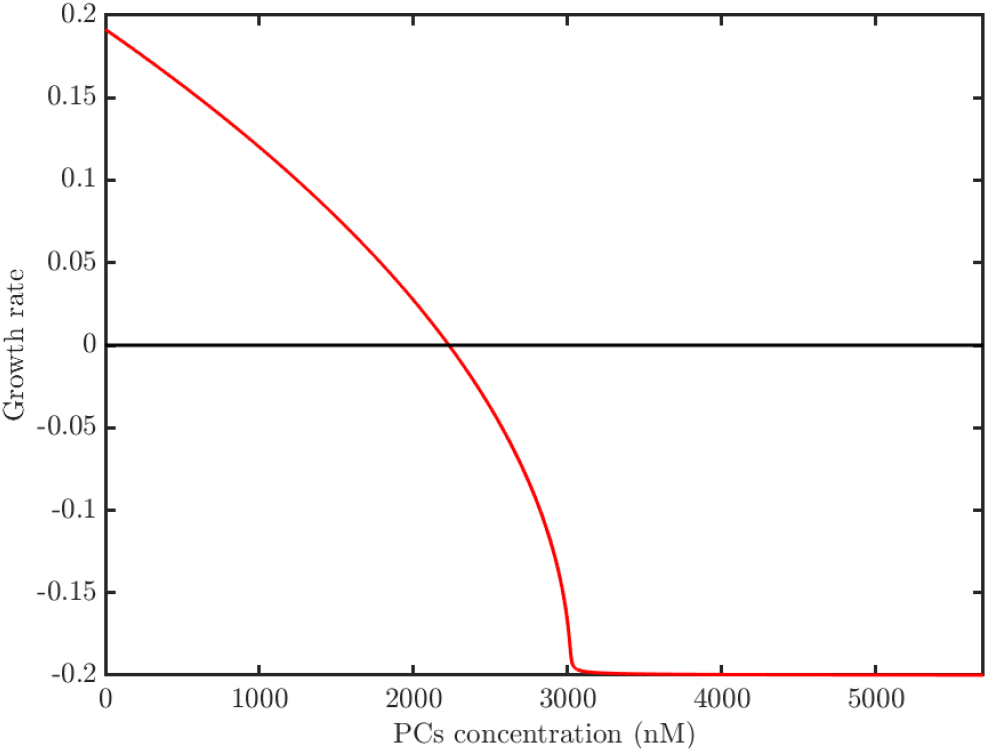
Effect of pharmacological chaperone dose on the prion growth rate

### Appendix D. 2. Data for antibodies

Since the available amount of PrPC protein drives the formation of nascent infectious proteins, reagents specifically binding prion-protein conformer may interrupt prion production. According to the experiments performed by Peretz et. al. [43] Fab D13 reduced the level of PrPSc compared with that in non-treated cells and it’s value for 50% inhibitory concentration (IC50) was 12*nM*. In addition, the levels of PrPC and glyceraldehyde-3-phosphate dehydrogenase in antibody-treated and untreated cells were found to be invariant, indicating that the prion protein antibodies used produced no cytotoxic effects that could have affected the PdPSc production. According to Regina R. Reimann [44], statistical analysis revealed significant lesion induction at 6 and 12*μ*g of D13, when those concentrations were injected, a conspicuous hyperintense lesion became apparent at 48*h*. To estimate the upper limit of the D13 intracerebrally injected safe dose, they performed a benchmark dose analysis which yielded a dose of 3.7–5.4*μ*g.

Moreover, the D13 antibody is lgG1 type antibody the half-life of these type of antibodies is dependent on the concentration, of approximately 29.7 days [36].

## Footnotes

1 One gene can form proteins that differ in both structure and composition; these different expressions are called isoforms [7]

2 A heterodimer is protein composed of non-identical monomers [9].

3 Pharmacological chaperones are small, cell-permeable molecules that assist correct protein folding [22].

## Notes

### Competing Interest Statement

The authors have declared no competing interest.

